# Uracil-DNA glycosylase efficiency is modulated by substrate rigidity

**DOI:** 10.1101/2022.08.30.505906

**Authors:** Paul B. Orndorff, Souvik Poddar, Aerial M. Owens, Nikita Kumari, Bryan T. Ugaz, Samrat Amin, Wade D. Van Horn, Arjan van der Vaart, Marcia Levitus

**Author notes:** To whom correspondence should be addressed: **ML**, **AvdV**, **WVH**. The authors wish it to be known that, in their opinion, the first three authors should be regarded as joint First Authors.

## Abstract

Uracil DNA-glycosylase (UNG) is a base excision repair enzyme that removes the highly mutagenic uracil lesion from DNA by a base flipping mechanism. UNG excision efficiency depends on DNA sequence, yet the underlying principles that dictate UNG substrate specificity have remained elusive. Here, we show that UNG efficiency is dictated by the intrinsic local deformability of the substrate sequence around the uracil. UNG specificity constants (*k*_cat_/*K*_M_) and DNA flexibilities were measured for an engineered set of DNA substrates containing AUT, TUA, AUA, and TUA motifs. Time-resolved fluorescence spectroscopy, NMR imino proton exchange measurements, and molecular dynamics simulations of the bare DNA indicated significant differences in substrate flexibilities. A strong correlation between UNG efficiency and substrate flexibility was observed, with higher *k*_cat_/*K*_M_ values measured for more flexible strands. DNA bending and base flipping were observed in simulations, with more frequent uracil flipping observed for the more bendable sequences. Experiments show that bases immediately adjacent to the uracil are allosterically coupled and have the greatest impact on substrate flexibility and resultant UNG activity. The finding that substrate flexibility controls UNG efficiency has implications in diverse fields, including the genesis of mutation hotspots, molecular evolution, and understanding sequence preferences of emerging base editors.

## INTRODUCTION

Cellular repair pathways maintain genetic integrity against thousands of daily spontaneous DNA lesions. The base excision repair (BER) pathway identifies, removes, and repairs small, non-helix distorting lesions. BER is initiated by specialized DNA glycosylases that catalyze the excision of the damaged base (1,2). DNA sequence effects have been identified for many glycosylases and alkyl-transferases that repair base mismatches, uracil bases, alkylated bases, etc. (3–8). In this work, we focus on understanding how DNA sequence impacts the repair of uracil, a highly mutagenic and common lesion in DNA (9,10). Uracil in DNA arises from dUTP misincorporation in place of dTTP during replication, or from spontaneous cytosine deamination. Unrepaired cytosine deamination results in mutagenic U:G mismatches, causing G:C to A:T transition mutations (11). While misincorporated uracil is not directly miscoding, it is a source of cytotoxic abasic sites in the genome (12,13). Abasic sites can block DNA replication and transcription (14) and can be converted into single-strand breaks by AP endonucleases (15).

Uracil-DNA glycosylase (UNG, also known as UDG) recognizes spurious uracil lesions as part of the BER pathway. The human UNG crystal structure bound to uracil-containing DNA (PDB: 1EMH) shows the uracil extruded from the duplex into the enzyme active site (16). This flipped-out complex exposes the damaged base before removal (17,18). The uracil base excision repair rate is controlled by the glycosylase that excises the lesion (19). In *E. coli*, the efficiency of uracil removal by UNG depends on DNA sequence and can vary by more than 15-fold, depending on the DNA context (5,6,20,21). Some general rules have emerged to explain UNG’s sequence dependence; for example, a vicinal thymine base 3’ of uracil generally results in poor uracil removal, and substrates with high local GC content are generally poor UNG substrates (20,21). Preferential excision of uracil does not correlate with DNA duplex melting temperatures; for instance, substrates containing the uracil in a TUA context are better UNG substrates than sequences with uracil in an AUT context (5,6). The differences between TUA and AUT sequences are particularly remarkable and are reminiscent of the asymmetries observed in the mechanical properties of undamaged DNA. In particular, TA steps are particularly flexible (deformable) with regard to roll, slide, and twist, while AT steps are more rigid (22–24). Nevertheless, to date, the underlying principles and substrate features that appear to dictate UNG enzyme efficiency remain elusive, despite fundamental impacts in diverse fields, including cancer and evolution.

For undamaged DNA, base and step flexibilities strongly depend on sequence (22,25–29), and are directly linked to mechanical properties such as persistence lengths and torsional rigidities (30). As structural studies have identified, the first steps that lead to the formation of the catalytically active UNG-DNA complex require significant conformational changes in the DNA, including hydrogen bond breakage and loss of stabilizing base-stacking interactions. Because these interactions are determined by the mechanical properties of DNA, the intrinsic deformability of the region surrounding the damaged site is likely an important and unrecognized variable in understanding the efficiency of the earliest events in the uracil excision process. A previous study using two different DNA substrates hinted at a connection between substrate flexibility and UNG activity (31), but the results were insufficient to establish the molecular basis for substrate preference. In this work, we test the hypothesis that UNG activity is dictated by the intrinsic local deformability of the DNA sequence around the uracil. Specifically, we expect uracil nucleotides embedded in rigid DNA contexts to be excised less efficiently than those incorporated in more deformable contexts.

To illuminate the link between UNG repair efficiency and substrate flexibility, we determined specificity constants (*k_cat_/K*_m_) for a variety of substrates and correlated the outcomes with the results of biophysical experiments and MD simulations that probe DNA substrate flexibility. A fluorescent analog of adenine, 2-aminopurine (2AP), was used to probe base stacking dynamics in the isolated substrates. Results show an inverse correlation between base stacking and *k_cat_*/*K*_m_, indicating a relationship between UNG activity and substrate flexibility. MD simulations on these substrates show markedly different dynamics around the uracil. Persistence lengths calculated from simulations are higher for DNA sequences that show a high degree of stacking in the fluorescence experiments. The analysis of the MD trajectories indicates that more flexible sequences show more frequent spontaneous base flipping of the uracil base, and NMR determinations of imino proton exchange rates validate these observations. Additionally, the NMR data show significant thermodynamic coupling between the bases flanking the uracil in the isolated substrates, and coupling between 5’ and 3’ flanking positions to uracil was also observed in UNG enzyme assays with the same substrates. Taken together, these results establish a clear link between the fundamental nature of substrate flexibility and the resulting UNG catalytic efficiency.

## MATERIALS AND METHODS

### Enzymes, oligonucleotides, and reagents

Uracil-DNA Glycosylase (UNG, MW=25.7 kDa) was purchased from New England BioLabs (Catalog # M0280L) at a concentration of 5,000 U/mL. Free 2-aminopurine riboside was purchased from AstaTech (PA, USA). Oligonucleotides containing canonical bases, uracil, or 2-aminopurine (2AP) were purchased from IDT (IA, USA) as desalted oligonucleotides. All sequences are listed in Table 1. Oligonucleotides for fluorescence experiments (kinetic assays, time-resolved fluorescence, and fluorescence quantum yields) were 39 nt in length (Table 1). Oligonucleotides for NMR experiments were shortened to the central 13 nucleotides to facilitate resonance assignments while still ensuring duplex formation. All oligonucleotides were solubilized in 1 × PBS buffer (20 mM sodium phosphate, 100 mM NaCl, 50 μM EDTA, pH 7.0). Concentrations were determined from measured absorbances at 260 nm using extinction coefficients provided by IDT.

**Table 1.**
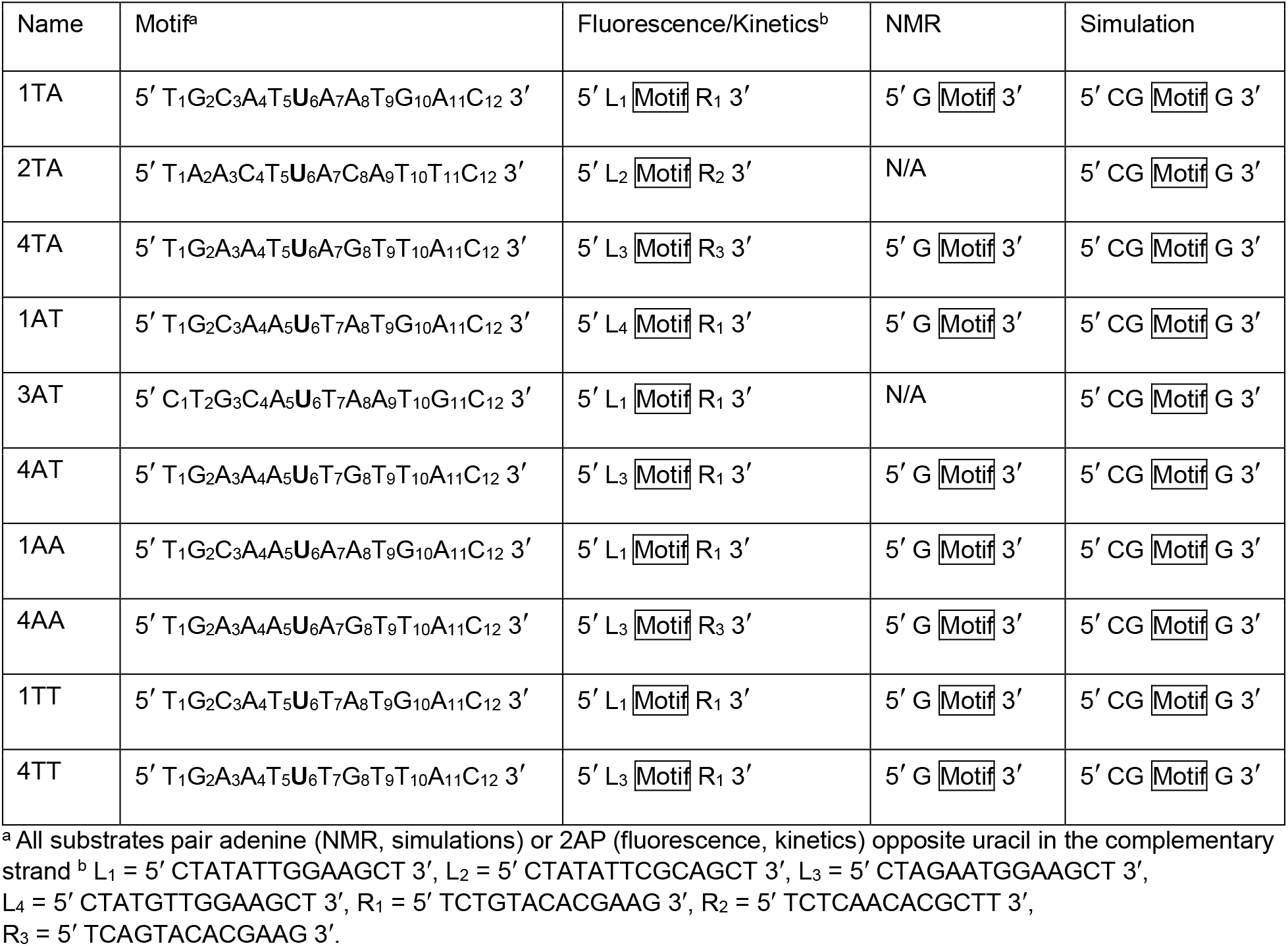
DNA Sequences.

### Sample preparation

Duplex DNA substrates used for the fluorescence-based experiments were prepared by annealing uracil-containing oligonucleotides with complementary strands containing 2AP opposite to the uracil. For the kinetic assays, a small excess of the 2AP-strand is preferable to an excess of the uracil-containing strand because the latter is also a substrate of UNG, and therefore its presence may affect the measured kinetic rates. For the kinetic assays, DNA substrates were prepared by annealing the strands at room temperature while monitoring the fluorescence intensity of 2AP in real-time. The uracil-containing strand was added to a known concentration of 2AP-containing strand, and the reduction in fluorescence intensity due to the formation of the duplex was measured until a small addition of the uracil strand did not result in a further decrease. The concentration of the resulting dsDNA substrate was calculated from the absorbance of the initial 2AP-containing strand and the volumes before and after adding the uracil strand. Traditional native polyacrylamide gel electrophoresis was used to confirm that the annealing procedure at room temperature was highly efficient and did not result in any measurable 2AP- or uracil-containing single strands. For the time-resolved and fluorescence quantum yield experiments, a slight excess of the U-strand is preferable to an excess of the 2AP-strand because 2AP in ssDNA is significantly brighter than 2AP in a duplex. Samples for these experiments were prepared as before and followed by the addition of a ~20% excess of the uracil-containing strand. In each case, we verified that further addition of the uracil-containing strand did not change the measured lifetimes or quantum yields. Duplexes for NMR experiments were prepared from complementary oligonucleotides mixed at 1:1 molar ratios that were heated to ~80 °C and annealed by cooling to room temperature for ~2 hours.

### UNG kinetic assay and data analysis

A continuous fluorescence kinetic assay was used to measure UNG activity on DNA substrates containing 2AP opposite to the uracil (32). 2AP is highly fluorescent when exposed to water but is highly quenched in dsDNA. The cleavage of the dU glycosidic bond by UNG results in an aldehydic abasic site opposite to 2AP, and this environmental change leads to a large fluorescence increase. The increase in fluorescence intensity can be used to calculate the reaction rate in a continuous kinetic assay. The kinetic parameters measured in this way are indistinguishable from the values obtained using a historical radioactivity-based electrophoresis assay with non-fluorescent substrates (32). 600 μL of duplex DNA was placed in a quartz micro cuvette (optical path length 10 × 2 mm) and fluorescence intensity (λ_ex_ = 310 nm, λ_em_ = 370 nm) was monitored continuously before and after addition of UNG. Fluorescence intensities were corrected in real-time for potential fluctuations in the incident intensity over the long measurement times. A small amount of stock enzyme (5,000 U/mL) was diluted 40-fold in 1× PBS buffer containing 1 mg/mL BSA, and this dilution was stored at 4 °C for no longer than six hours and used for all kinetic experiments performed within the same day. For the kinetic experiments, 2 μL of this dilution were added to the cuvette containing the DNA substrate for a final concentration of UNG in the assay mixture of 0.16 nM. The initial velocity (V_0_, units of M·s^−1^) was obtained from the initial slope of the measured intensity (*F*(*t*)) as 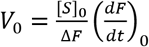, where [S]_0_ is the concentration of DNA substrate (0.075-6 μM) and ΔF is the total change in fluorescence. A representative sample run and a sample calculation are shown in Fig. S1. Error bars in Figs. 1 and S7 represent 95% confidence intervals. To rule out systematic sources of error, for 1AT, 23 trials were performed at 0.3 μM concentration involving 1) at least four different purchased UNG stock solutions, 2) three different experimentalists, and 3) DNA substrates prepared from oligos purchased at different times. The V_0_ vs [DNA] curves were fitted using the Michaelis-Menten equation in Origin Pro (Northampton, MA) using the Lavenberg Marquardt iteration algorithm and using the reciprocal of the variances as weights.

**Figure 1.**
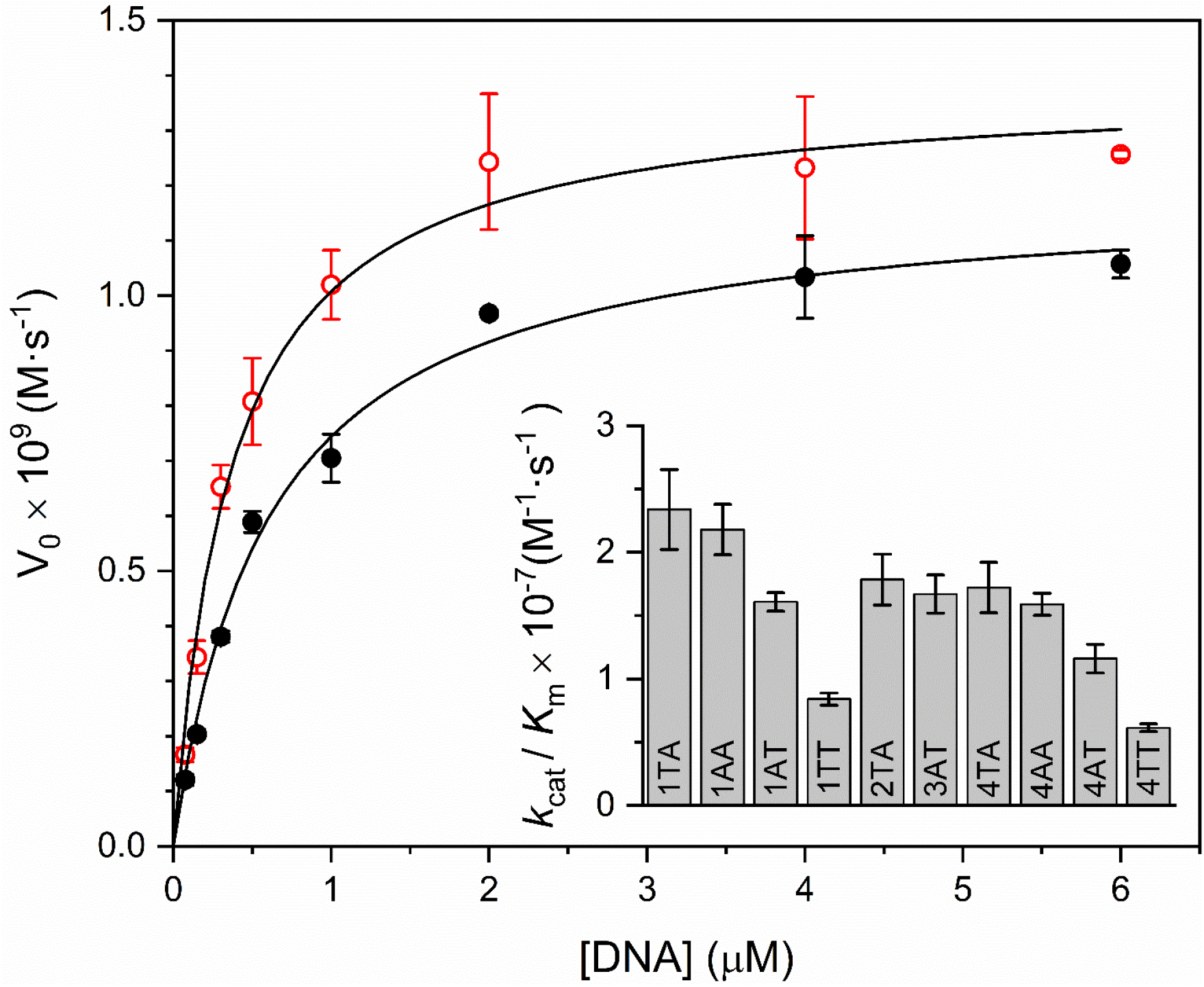
Experimental initial rates (V_0_) against substrate concentration for substrates 1TA (red open circles) and 4AT (black solid circles). Results for all other samples are shown in Fig. S7. Each data point is the mean of at least three independent experiments. Error bars represent 95% confidence intervals. The data have been fitted to the Michaelis-Menten equation (see table S2 for all Michalis-Menten parameters). Inset: Values of *k*_cat_/*K*_M_ for all substrates (see Table S2).

### UNG assays with competing substrates

Kinetic assays were carried out with two competing substrates in solution to obtain relative catalytic efficiencies. Two experiments were performed for each pair of substrates. In the first, substrate *x* contained 2AP opposite to the uracil, while substrate *y* contained a canonical adenine. The velocity measured in this experiment is *V_x_*, the rate of uracil removal from substrate *x* when the enzyme competes for substrates *x* and *y*. The second experiment was carried out with 2AP-labeled substrate *y* and unlabeled *x*, from which *V_y_* was calculated. The ratio *V_x_/ V_y_* equals the ratio of the catalytic efficiencies, 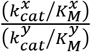, when the concentrations of substrates *x* and *y* are equal and kept constant in both experiments (33,34). Competition experiments were carried out for three pairs of substrates (1TA/1AT, 4TA/4AT, and 1TA/4TA, see table 1) with solutions containing 0.3 μM of each substrate and 0.16 nM UNG.

### Time-Resolved Fluorescence

Time-resolved fluorescence intensity measurements were performed using the time-correlated single-photon counting (TCSPC) technique. A mode-locked Ti:Sapphire laser (Mira 900, Coherent) pumped by a frequency-doubled Nd:YVO4 laser (44% from an 18 W Verdi, Coherent) was used as the excitation source. The 130 fs light pulses (at 800 nm with a repetition rate of 250 KHz) were generated by a regeneratively amplified Ti:S laser system (RegA 9000, Coherent Laser). The pulses were sent to an optical parametric amplifier (OPA) to generate the excitation light at 620 nm and then frequency-doubled to obtain excitation pulses at 310 nm. Fluorescence emission was collected at a 90° geometry setting and detected using a double-grating monochromator (Oriel Instruments) and a microchannel plate photomultiplier tube (Hamamatsu R3809U-51). Decays were measured at three emission wavelengths (380, 390, and 400 nm) for global analysis as described below. The polarization of the emission was 54.7° relative to that of the excitation (magic angle). A single-photon counting card (Becker-Hickel, SPC-830) using two time windows (3.3 ns and 25 ns) was used for data acquisition. Instrumental response functions (IRF) were determined for both time resolutions. The typical IRF had a FWHM of 40 ps, measured from the scattering of Ludox sample at 310 nm. The three decays obtained at different emission wavelengths for each sample were fitted globally keeping the lifetimes as common parameters among the three data sets. This approach minimizes the problem of correlation between pre-exponential factors and lifetimes, which is common when fitting multi-exponential decays (35). The fitting parameters were obtained through iterative reconvolution of the model function 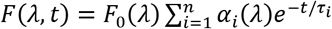 with the measured IRF using an in-home written software package (ASUFIT). Here, λ represents the emission wavelength, and 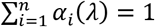. Lifetimes as long as ~8 ns and as short as ~30 ps are expected (35), and in our experience, results are more robust and reproducible if the shorter components are determined from data measured with the highest time resolution achievable with the single-photon counting card (814 fs/channel, 2^12^ channels = 3.3 ns total), while the longer lifetimes are obtained from data measured with a wider time window. The decays measured using 3.3 ns acquisition windows were fitted using three exponential terms. A fourth term did not improve the quality of the fit. A new fit was then conducted using the decays measured with a 25 ns acquisition window (6.1 ps/channel) using four exponential terms but fixing the two shortest lifetimes (τ_1_ and τ_2_) to the values obtained in the previous fit. In this way, the fitting parameters in the second fit were τ_3_, τ_4_ and α_1–4_(*λ*). This procedure allows a more accurate determination of τ_1_ and τ_2_ (both below 0.5 ns) from measurements using 814 fs/channel resolution and τ_3_ and τ_4_ (both over 1 ns) from measurements using 6.1 ps/channel resolution. Mean lifetimes were calculated for each sample as 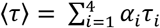 using the *α_i_* values obtained in the second fit.

### Fluorescence quantum yields

Fluorescence quantum yields (ϕ) were determined relative to a reference as 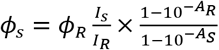, where the subscripts “S” and “R” refer to the sample and reference, respectively, *I* is the integrated emission intensity measured over the entire fluorescence band, and A is the absorbance at the excitation wavelength (315 nm) (36). Absorbances were kept below 0.05 to avoid inner filter effects. Experiments replacing the 2AP-containing strand with an adenine-containing strand were used to verify that the absorbance of the canonical bases at 315 nm is negligible compared to the absorbance measured with the 2AP-containing DNA. Steady-state emission fluorescence spectra were acquired on a PTI Quantamaster 4/2005SE spectrofluorimeter. Fluorescence spectra showed clear contributions from Raman scattering at 352 nm, and to account for these contributions a buffer sample was used as a blank and subtracted from the measurements performed with 2AP-containing DNA. The free 2AP riboside is commonly used as a reference (*ϕ_R_* = 0.68 in water) (37), but fluorescence intensities in duplex DNA are reduced 100-fold or more due to quenching, and therefore the determination of ϕ in DNA involves measuring very small fluorescence intensities. To improve accuracy, we performed five independent determinations of ϕ for the sample with the highest quantum yield (1AT) using free 2AP in water as a reference. The average of five determinations was ϕ_1AT_ = 0.0103 (standard deviation = 6.5 × 10^−4^, 95% confidence interval = 8.1 × 10^−4^), and this value was subsequently used as a reference for the ϕ determinations of all other 2AP-DNA samples. Values listed in Table S3 are averages of 4 independent determinations. All standard deviations are 3% or lower.

### Fractional population of highly stacked species (α_0_)

The fractional population of dark (highly stacked) 2AP molecules in the duplex DNA substrates was calculated from the sample mean lifetime (〈*τ*〉) and fluorescence quantum yield (ϕ) as 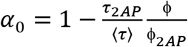, with *ϕ_2AP_* = 0.68 and *τ_2AP_* = 10.2 ns (37,38).

### NMR-detected Imino Proton Exchange Rate Measurements

NMR samples were prepared in 3 mm tubes with DNA concentrations ranging from 2 mM to 4 mM with 5% v/v deuterium oxide. NMR experiments were performed on a Bruker 850 MHz Avance III HD spectrometer equipped with a 5 mm TCI CryoProbe, and a Bruker 600 MHz Avance III HD spectrometer equipped with a Prodigy probe. NMR spectra were processed and analyzed using Bruker TopSpin 4.1, MestReNova 14.2, and Matlab 2019b.

#### Resonance assignments

Two-dimensional ^1^H, ^1^H – Nuclear Overhauser Effect Spectroscopy (NOESY) experiments were recorded at 20 °C utilizing water suppression by excitation sculpting. The resulting 2D spectra were used to assign imino protons for each duplex using traditional “backbone-walking” methods (Figs. S2-S3) (39,40). All assignments and future experimentation were collected at 20 °C.

#### Water inversion efficiency factor (E)

The water inversion efficiency factor was measured as previously described (41), with a relaxation delay of 30 s, and is further described in the supplementary information (Fig. S4). Data processing and fitting were completed in Bruker TopSpin 4.1 and MestReNova 14.2.

#### Longitudinal relaxation rate of water (R_1w_)

The relaxation rate of water (*R_1w_*) was measured utilizing a previously described saturation-recovery method that is compatible with high Q cryoprobes (41). Variable time delays ranged from 1 ms to 18 s. Data processing and fitting were completed in Bruker TopSpin 4.1 and Matlab 2019b for verification (Fig. S5). Determination of *R_1w_* was completed using the TopSpin T1/T2 Module.

#### Imino proton longitudinal relaxation (R_1n_) and exchange rates (k_ex_)

The sum of the longitudinal imino proton relaxation rates (*R_1_*) and the imino proton solvent exchange rate (*k_ex_*) can be used to determine the longitudinal relaxation rate of each imino proton (*R_1n_*). A pseudo-two-dimensional experiment was implemented to determine the longitudinal relaxation of imino protons (*R_1n_*) and the imino proton exchange rates (*k_ex_*) following established methods (42) utilizing a 24-point variable delay sequence ranging from 1 ms to 15 s. The respective spectra were processed in TopSpin 4.1, and the data were fit with MATLAB R2019b (MathWorks) using nonlinear least-squares fit, first fitting for the longitudinal relaxation of imino protons (*R_1n_*) followed by the exchange rate of imino protons (*k_ex_*). Representative fits of *R*_1n_ and *k_ex_* are shown in Fig. S6. The *k_ex_* was determined by fitting the individual peak areas to the equation:

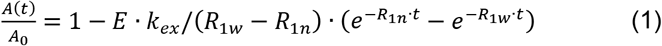

where *A*(*t*) is the area of the peak at exchange time point *t, A_0_* is the peak intensity at equilibrium, *E* (determined experimentally) is the water inversion efficiency factor, *R_1w_* (determined experimentally) is the water longitudinal relaxation rate, and *R_1n_* (determined experimentally) is the sum of imino proton longitudinal relaxation rates and its solvent exchange rate. The reported errors were estimated from the fitting of *k_ex_* (Table S1).

#### Double mutant cycles and base-pair coupling analyses

The energetic impact of base substitutions was quantified from thermodynamic cycles that depict differences in transition state free energies: 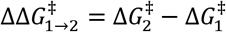, where indices 1 and 2 indicate two different DNA sequences (43). Two cycles were constructed. The first was based on NMR data, where Δ*G*^‡^ represents the barrier for imino exchange, and 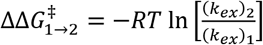. Differences between the 4AT, 4TT, 4AA, and 4TA sequences were assessed. The second cycle was based on *k*_cat_/K_m_ measurements, where Δ*G*^‡^ represents the barrier for enzymatic uracil excision, and 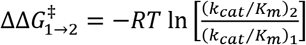. In this cycle, differences between 4AT, 4TT, 4AA, 4TA, and 1AT, 1TT, 1AA, 1TA (see Table 1 for substrate nomenclature) were calculated. From these cycles, coupling free energies

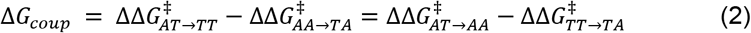

between the base pairs directly adjacent to uracil were assessed. Δ*G_coup_* is zero when these base pairs are independent of each other, and nonzero when they are coupled and influence each other. Since the transition state free energies of the two cycles correspond to different processes, free energy differences and coupling free energies are expected to differ between the thermodynamic cycles.

### MD simulations

The dsDNA sequences of Table 1 were built in the unbent BII conformation using 3DNA (44); U_6_ was base-paired to A_19_. Each strand was solvated in a rectangular TIP3P (45) water box of 100 mM NaCl with a solvent layer of 15 Å in each direction. After energy minimizations, each system was heated from 100 to 300 K over 2.5 ns with a 1 kcal/(mol·Å^2^) harmonic restraint on all DNA atoms. During heating, flat bottom distance restraints with a force constant of 1 kcal/(mol·Å^2^) were added to the hydrogen bonds between the bases. After heating, the harmonic restraints on the DNA atoms were gradually removed over 1.2 ns, while restraints on the hydrogen bonds remained in effect. The latter were subsequently removed over an additional 3 ns. The unrestrained systems were then equilibrated for 400 ns, followed by at least 600 ns of production simulations. Heating and restraints removal were performed with Langevin dynamics in AMBER (46), while the production runs were done with Langevin dynamics in OPENMM (47). The simulations were performed in NPT, periodic boundary conditions were in effect, SHAKE (48) was applied to all covalently bonded hydrogen atoms, and long-range electrostatic interactions were handled using the particle-mesh Ewald method (49). All simulations used the AMBER OL15 DNA force field (50); deoxyribose parameters for U were taken from T deoxyribose. Convergence was assessed by monitoring cumulative averages of DNA bending and total winding angles. In addition, all trajectories were decorrelated using pymbar (51), and all properties were calculated from 100 decorrelated frames per trajectory. If needed, simulations were extended for additional blocks of 500 ns until convergence. Simulations were run in triplicate for each strand.

Geometric analyses of the DNA step parameters were performed with 3DNA (44). DNA bending angles (*ϕ*) were calculated from tilt, roll, and twist base step angles using the MADBEND procedure (52,53). Bending persistence lengths (BPLs) were calculated from: 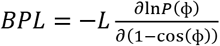 (54), where *L* is the contour length, and *P*(*ϕ*) the probability of observing a particular bending angle. Contour lengths were calculated from the sum of the helical rise; to account for fraying, the terminal base steps were excluded from the *ϕ* and BPL analyses. Torsional persistence lengths (TPLs) were calculated from the variance of the total winding angle 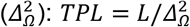 (55). The total winding angle was calculated as the sum of the individual twist steps for each sequence. To account for fraying and base flipping, the two terminal base steps and those neighboring U_6_ were excluded from the sum.

Extrahelical flipping of uracil was assessed by monitoring the hydrogen bonding distance between HN_3_ of U_6_ and N_1_ of its complementary A_19_, and the flipping angle. This flipping angle was taken as the pseudo-dihedral angle between the center of mass (COM) of the base ring of U_6_, the COM of the U_6_ backbone, the COM of the backbone of residue 8, and the COM of the backbone of the base complementary to U_6_ (56). Based on visual inspections of the trajectories, U_6_ was considered flipped out when the U_6_ - A_19_ hydrogen bond distance exceeded 4.5 Å and the pseudo-dihedral angle was greater than 40 or less than −40 degrees. Negative pseudo-dihedral angles correspond to major groove flipping, while positive angles correspond to minor groove flipping.

## RESULTS

### DNA Substrates

A previous study reported relative excision efficiencies for uracils embedded in different sequence contexts within a long (> 6 kbp) viral dsDNA genome (6). Inspection of the DNA sequences reported in this study suggests that AUT sequences are generally poorer UNG substrates than TUA sequences despite similar overall AT/GC content (6). Furthermore, there are clear variations within each group, indicating that bases not immediately 3’ or 5’ of the uracil also contribute to UNG activity. Based on these published data, we designed a number of dsDNA substrates containing uracil in TUA or AUT contexts, and adenine opposite uracil (Table 1). Substrates 1TA, 2TA, 3AT, and 4AT in Table 1 share the same central motif as sequences reported in ref. 6 and were designed to span a range of uracil excision efficiencies. Relative percent uracil removal values for these sequences embedded in a viral dsDNA genome were reported to be 100%, (50 ± 13)%, (35 ± 7)%, and (12 ± 7)%, respectively (6). Substrates 1AT and 4TA were additionally designed to evaluate the impact of swapping the flanking A and T bases of the substrates with the best (1TA) and worst (4AT) uracil removal efficiency in this list. Lastly, selected substrates containing uracil in a AUA (1AA and 4AA) or a TUT (1TT and 4TT) context were designed to evaluate the contributions of nucleotides flanking the uracil.

### Kinetics of uracil removal by UNG and competition assays

A continuous fluorescence kinetic assay was used to measure UNG activity on DNA substrates containing 2AP opposite to the uracil (32). An example of a kinetic run and initial velocity calculation is included in the supplemental materials (Fig. S1). Initial reaction velocities (V_0_) were obtained from the early stages of the reaction progress curves, and Michaelis-Menten parameters (*K*_m_ and *k*_cat_) were obtained from a nonlinear fit of the experimental V_0_ vs. [DNA] data (Figs. 1 and S7). Michaelis-Menten parameters and specificity constants (*k_cat_*/*K*_m_) for the different substrates are listed in Table S2. For substrates 1TA, 2TA, 3AT, and 4AT, calculated *k_cat_*/*K*_m_ ratios parallel the percent uracil removal efficiencies reported previously, albeit in longer and more complex DNA substrates: 1TA > 2TA > 3AT > 4 AT (Fig. 1, inset). For substrates 1AT and 4TA, which are identical to 1TA and 4AT except for the swapped flanking bases, *k_cat_*/*K*_m_ ratios are higher in a TUA context compared to AUT (i.e. 1TA > 1AT and 4TA > 4AT, Fig. 1 and Table S2).

Competition experiments with UNG acting on two different uracil-containing substrates present in the same reaction mixture were additionally performed for three pairs of substrates: 1TA/1AT, 4TA/4AT, and 1TA/4TA. Values of 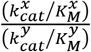 obtained in this way are *V*_1*TA*_/*V*_1*AT*_ = 1.65, *V*_4*TA*_/*V*_4*AT*_ = 1.59, and *V*_1*TA*_/*V*_4*TA*_ = 1.44. These values agree with the 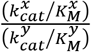 ratios calculated from the *k*_cat_ and *K_m_* values obtained from the Michaelis-Menten fits within experimental error. The first two pairs (1TA/1AT and 4TA/4AT) confirm that UNG’s specificity is greater for uracil in a TUA context compared to AUT. The third pair (1TA/4TA) indicates that substrate specificity is not solely determined by the flanking bases (see also Fig.1, inset).

### Fluorescence quantum yields and time-resolved fluorescence

2AP is an isomer of adenine (6-aminopurine) that, unlike the canonical DNA bases, is highly fluorescent when free in solution. The fluorescence quantum yield of 2AP in water is 0.68 (37), and its fluorescence intensity decay is monoexponential with a single lifetime of 10-11 ns (57). When incorporated into DNA, 2AP fluorescence is quenched strongly by interactions with its neighboring bases (35,58–61). Inter-base interactions in duplex DNA give rise to a multiexponential decay that reflects the highly heterogeneous environment sensed by the probe (38,60–63). We used steady-state and time-resolved 2AP fluorescence to probe DNA dynamics around the uracil lesion. Consistent with previous reports (60,61,63), four exponential terms with lifetimes ranging from tens of picoseconds to nanoseconds were needed to fit the time-resolved TCSPC data (Table S4):

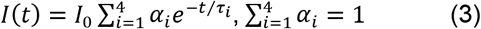

Lifetimes in the picosecond timescale have been reported for 2AP in dsDNA using ultrafast methods (64), indicating that a fraction of the emitting 2AP molecules has ultrafast lifetimes that fall outside the resolution of the TCSPC technique (~40 ps) (38). The fractional population of the highly stacked probes that give rise to lifetimes below the resolution of the measurement (denoted by α_0_) was determined from the average lifetimes (Table S4) and the measured fluorescence quantum yields (Table S3), as described in Materials and Methods (38,65). Calculated α_0_ values are shown in Table S5 and Fig. 2.

**Figure 2.**
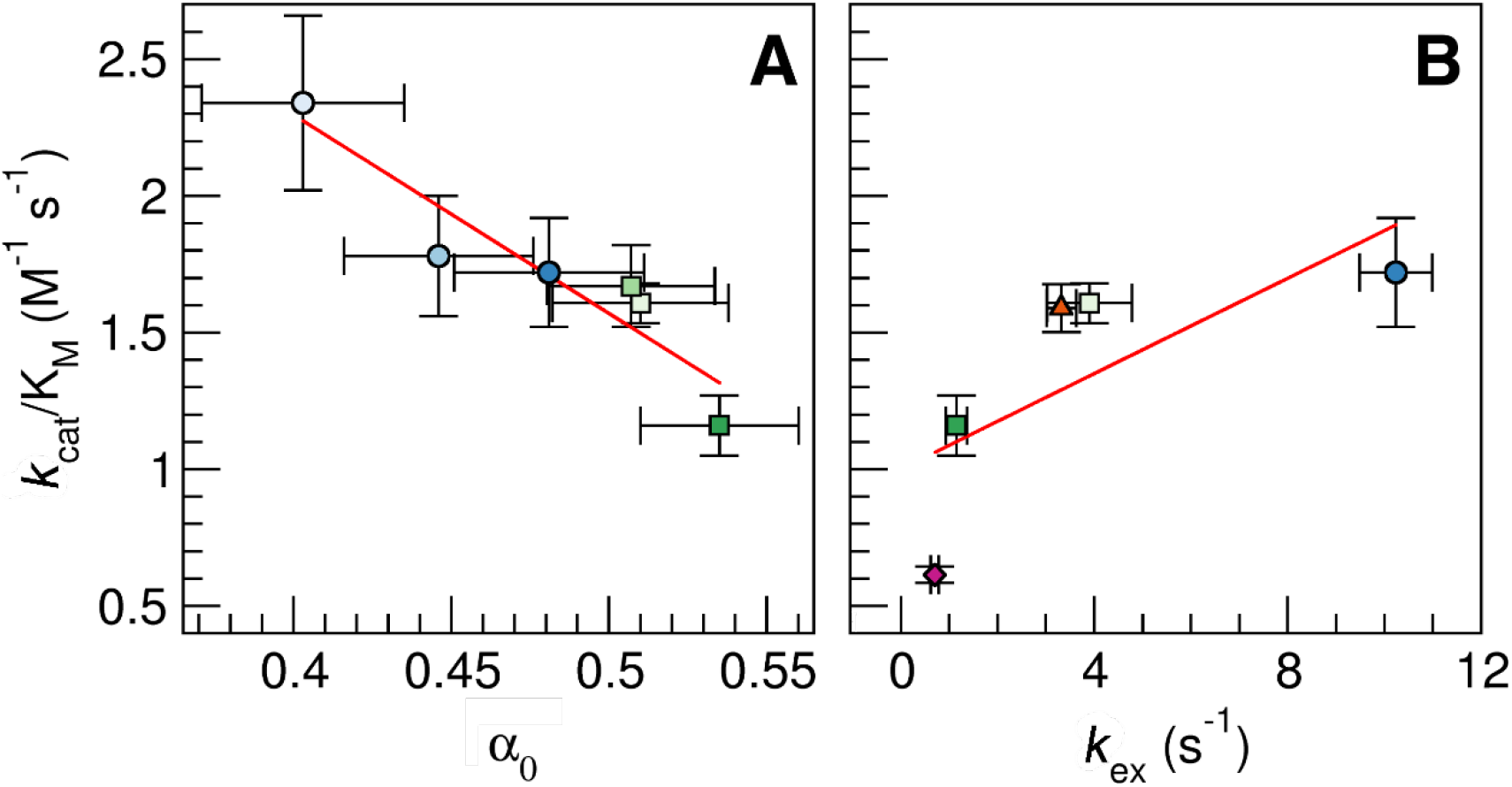
Correlation between *k*_cat_/*K*_M_ values and α_0_ (A) and *k*_ex_ of U_6_ (B). Red lines indicate linear regressions; with correlation coefficients of −0.930 (A) and 0.727 (B), respectively. 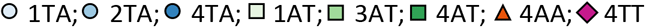.

Seibert et al. reported measurements of fluorescence quantum yields and lifetimes of 2AP placed opposite to a uracil in two different 19 bp DNA substrates: one containing uracil flanked by two As (AUA, high UNG efficiency) and one containing uracil flanked by two Gs (GUG, low UNG efficiency).(31) A shorter average lifetime (〈*τ*〉) was measured for AUA (0.32 ns) than for GUG (2.48 ns), and this was interpreted as indicative of AUA being more flexible than GUG, leading to more efficient dynamic quenching of the former. Guanine, however, is an efficient quencher of 2AP (59,66), and 2AP lifetimes as low as 400 fs, were measured for 2AP in the vicinity of G (64). The surprisingly long average lifetime reported for this sequence (2.48 ns), therefore, likely reflects the fact that most of the 2AP population is quenched dynamically with lifetimes shorter than the resolution of the experiment (the shorter lifetimes reported were ~200 ps, which are relatively high for TCSPC measurements). Had the authors measured the short lifetimes with high relative amplitudes expected for 2AP in the vicinity of G, the calculated 〈*τ*〉 would have been significantly shorter and likely below the value measured for AUA. These arguments illustrate the problems with interpreting relative 2AP average lifetimes in terms of substrate flexibility. Quenching by the different DNA bases may occur by different mechanisms and in different timescales, and measured average lifetimes are sensitive to instrumental resolution (35). To avoid these pitfalls, we favor the currently accepted view that the relative contributions of each lifetime (α_i_), but not the lifetimes themselves, are useful measures of the degree of base stacking, and therefore reflect substrate flexibility (35).

The excited 2AP population is expected to be partitioned between several different local environments that lead to different quenching efficiencies. Each lifetime represents a distribution of conformations in which 2AP experiences similar quenching rates. The four lifetimes commonly found in TCSPC measurements (30-50 ps resolution) include values from ~8 ns to less than 100 ps, representing increasingly stacked conformations (35). The corresponding normalized amplitudes, α_i_ (Eq. 3), measure the fractional population of each conformation detectable by TCSPC, from more stacked (α_1_), to more exposed to the solvated environment (α_4_) (35). In addition, a fifth population (α_0_) of highly stacked 2AP molecules results in ultrafast quenching and ultrashort lifetimes that TCSPC does not detect. The existence of this population is evident from the fact that fluorescence quantum yields and fluorescence lifetimes are not proportional to each other. For example, 〈*τ*〉 is greater for 4TA than for 4AT, but the opposite is true for the fluorescence quantum yields (ϕ_4AT_ > ϕ_4TA_). This indicates that the fractional population of 2AP molecules with lifetimes shorter than ~40 ps (TCSPC resolution) is greater in 4AT than in 4TA. All calculated α_0_ values are quite high (α_0_ > 0.4 for all sequences, Table S5), but values are higher for substrates containing uracil in a AUT context compared to TUA. A higher α_0_ value indicates a higher fraction of highly stacked (and therefore efficiently quenched) 2AP molecules, which we interpret as evidence for a less deformable substrate.

Substrates 1AT and 4TA were designed from parent substrates with high (1TA) and low (4AT) UNG efficiency to test the hypothesis that differences in substrate deformability determine preferential repair efficiencies. Swapping the flanking A and T while keeping the rest of the sequence constant affects stacking interactions in the vicinity of the uracil without changing the substrate melting temperature. This swap resulted in a higher fraction of highly stacked 2AP probes for substrates containing the uracil in a TUA/TPA context (where P denotes 2AP, which substitutes for A), i.e α_0_ (1AT) > α_0_ (1TA) and α_0_ (4AT) > α_0_ (4TA). As noted above, *k*_cat_/*K*_m_ values follow the opposite trend, suggesting a correlation between substrate deformability and uracil removal efficiency by UNG. A strongly negative correlation between *k*_cat_/*K*_m_ and α_0_ was indeed observed for all sequences within experimental uncertainty (Fig. 2A), indicating that more flexible sequences have higher repair rates.

### Molecular Dynamics Simulations

We performed MD simulations to quantify the flexibility of the various DNA duplexes (Table 1), and to gain molecular insights into the origins of the differences observed in the time-resolved fluorescence experiments. All simulations were repeated in triplicate. Calculated bending persistence lengths showed that all TA sequences and 3AT were more flexible than undamaged DNA (which has a persistence length of ~500 Å) (67,68), while the 1AT, 4AT, 1TT, 4TT, 1AA, and 4AA sequences were similar to undamaged DNA in bending rigidity (Table S6). We observed a clear distinction between the TA and AT sequences, with the TA sequences having lower bending persistence lengths than the AT sequences. The AA and TT sequences were least flexible in terms of bending. The higher flexibility of the TA sequences was largely echoed in calculated torsional persistence lengths: the AT sequences generally had larger torsional persistence lengths than the TA sequences. The only exception was the 3AT sequence, which was more flexible than 4TA, the stiffest TA sequence. Torsional persistence lengths of the AA and TT sequences were similar to the AT sequences. The standard deviation of the bending angle showed clear distinctions between the TA, AT, and AA/TT sequences. It was highest for the TA sequences, indicating large bending flexibility of these sequences, and lowest for AA/TT, indicating higher rigidity (Table S6). Overall, calculated properties indicated the highest flexibility for the TA sequences and the lowest flexibility for the AA and TT sequences. Fig. 3 shows that these three properties are strongly correlated with the α_0_ values obtained from fluorescence measurements. The correlation coefficient (*r*) was 0.918 for the bending persistence length, 0.869 for the torsional persistence length, and −0.939 for the standard deviation of the bending angle. Moreover, ranking of the sequences by flexibility was largely similar for these MD measures and α_0_.

**Figure 3:**
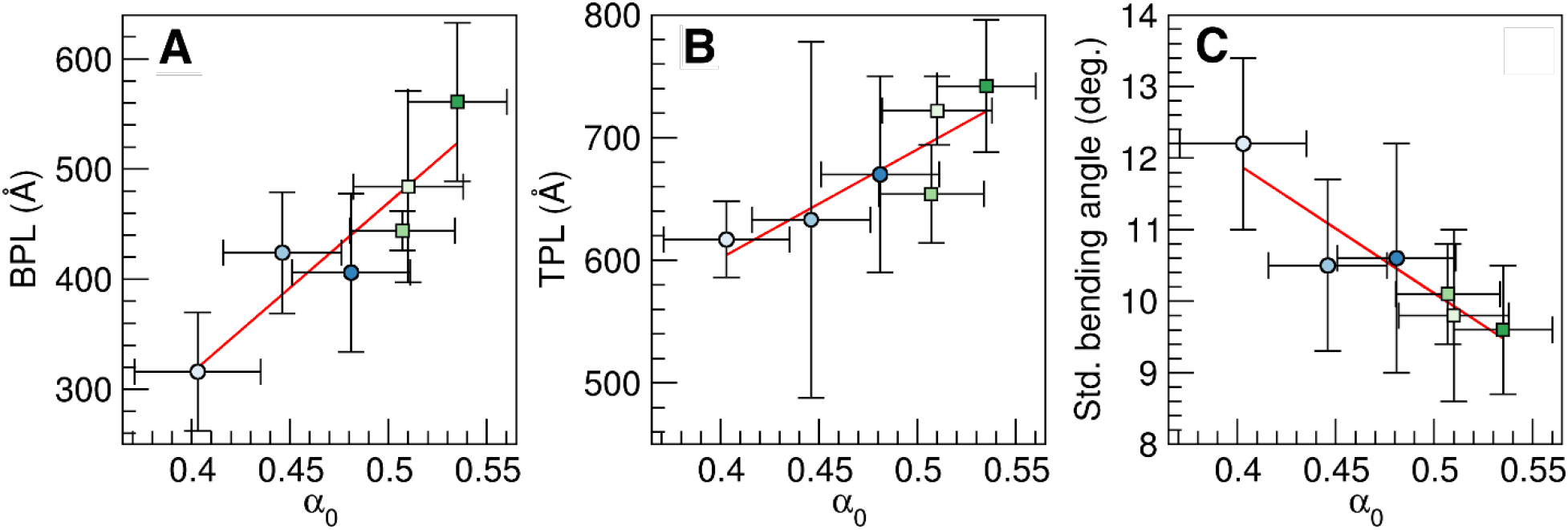
Correlation of MD properties with α_0_. A) Bending persistence length (in Å), B) torsional persistence length (in Å), C) standard deviation of the bending angle (in degrees). Averages and standard deviations over 3 MD replicas. Red lines show linear regression of the data; correlation coefficients are 0.981 (A), 0.869 (B), and −0.939 (C). 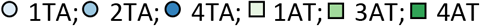.

The sequences displayed markedly different local dynamics around the lesion. Fig. S8 shows the shift, slide, and rise translational step parameters, and tilt, roll, and twist rotational parameters for the central A_5_U_6_A_7_, A_5_U_6_T_7_, T_5_U_6_A_7_, and T_5_U_6_T_7_ steps. These values were averaged over all trajectories and all sequences; standard deviations, shown as bars, are a measure of the base step flexibilities. Interestingly, differences in UA and AU step flexibilities depend on context and do not completely mirror the behavior of the TA and AT steps of undamaged DNA (22–24). For example, UA is particularly flexible in the TUA motif, but rigid in the AUA motif. The TUA motif was by far most flexible. It displayed large flexibilities in all step parameters of both steps. We observed some asymmetries in the UA and TU steps of this motif. The roll angle of its UA step was more flexible than its TU step, its TU step was more flexible than UA in slide and twist, while both steps had similar flexibilities for the other parameters. Second most flexible was the AUT motif. Its UT step was more flexible in roll, twist, shift, and rise, its AU step was more flexible in slide, and its tilt flexibility was similar for both steps. In contrast, the AU, UA, TU and UT steps of the AUA and TUT motifs displayed low flexibilities.

The main reason for the high flexibility of the central base steps of the TUA sequences was extra-helical base flipping of U_6_ (Table 2). Extra-helical base flipping was observed in all TUA sequences. Flipping was reversible and would occur throughout the simulations, but in 4TA and 2TA U_6_ remained extra-helical for nearly the entirety of the simulations. Such spontaneous base flipping of uracil damaged DNA has also been observed in previous MD simulations (69). Base flipping was also observed in the AUT sequences, particularly in the 3AT and 1AT sequences, but this was by far not as prominent as when uracil was in a TUA context. 1AA and 4AA displayed even less flipping than the AUT sequences, and flipping did not occur in 1TT and 4TT.

**Table 2.**
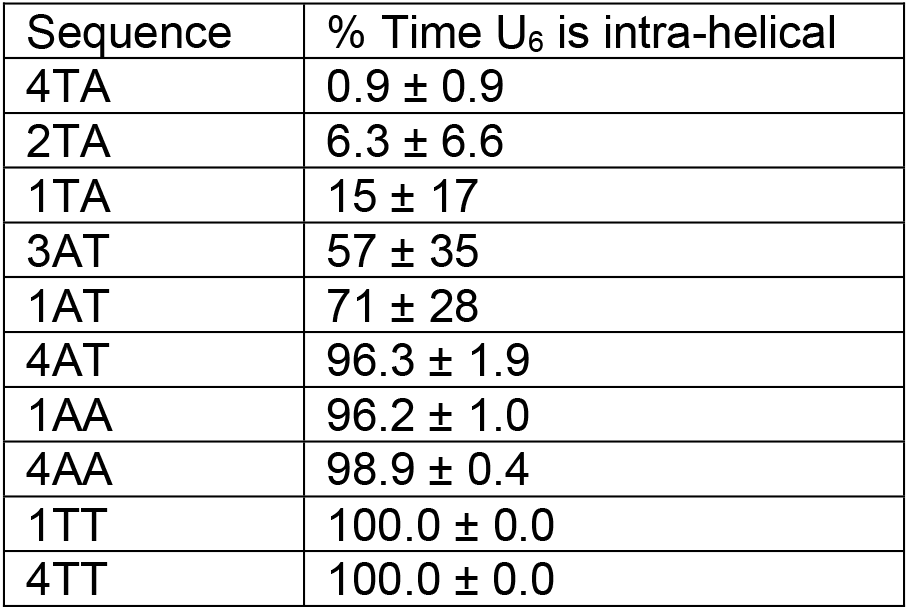
Extra-helical flipping of U_6_. Average and standard deviations over three MD replicas.

| Sequence | % Time U<sub>6</sub> is intra-helical |
| --- | --- |
| 4TA | 0.9 ± 0.9 |
| 2TA | 6.3 ± 6.6 |
| 1TA | 15 ± 17 |
| 3AT | 57 ± 35 |
| 1AT | 71 ± 28 |
| 4AT | 96.3 ± 1.9 |
| 1AA | 96.2 ± 1.0 |
| 4AA | 98.9 ± 0.4 |
| 1TT | 100.0 ± 0.0 |
| 4TT | 100.0 ± 0.0 |

Flipping of U_6_ was highly correlated to bending motions. A measure of the flipping motion is the standard deviation of the flipping angle. This standard deviation was highly correlated to the bending persistence length and the standard deviation of the bending angle, with correlation coefficients of −0.972 and 0.897, respectively (Fig. 4). The correlation to the torsional motion was weaker, with a correlation coefficient of −0.763 for the torsional persistence length.

**Figure 4:**
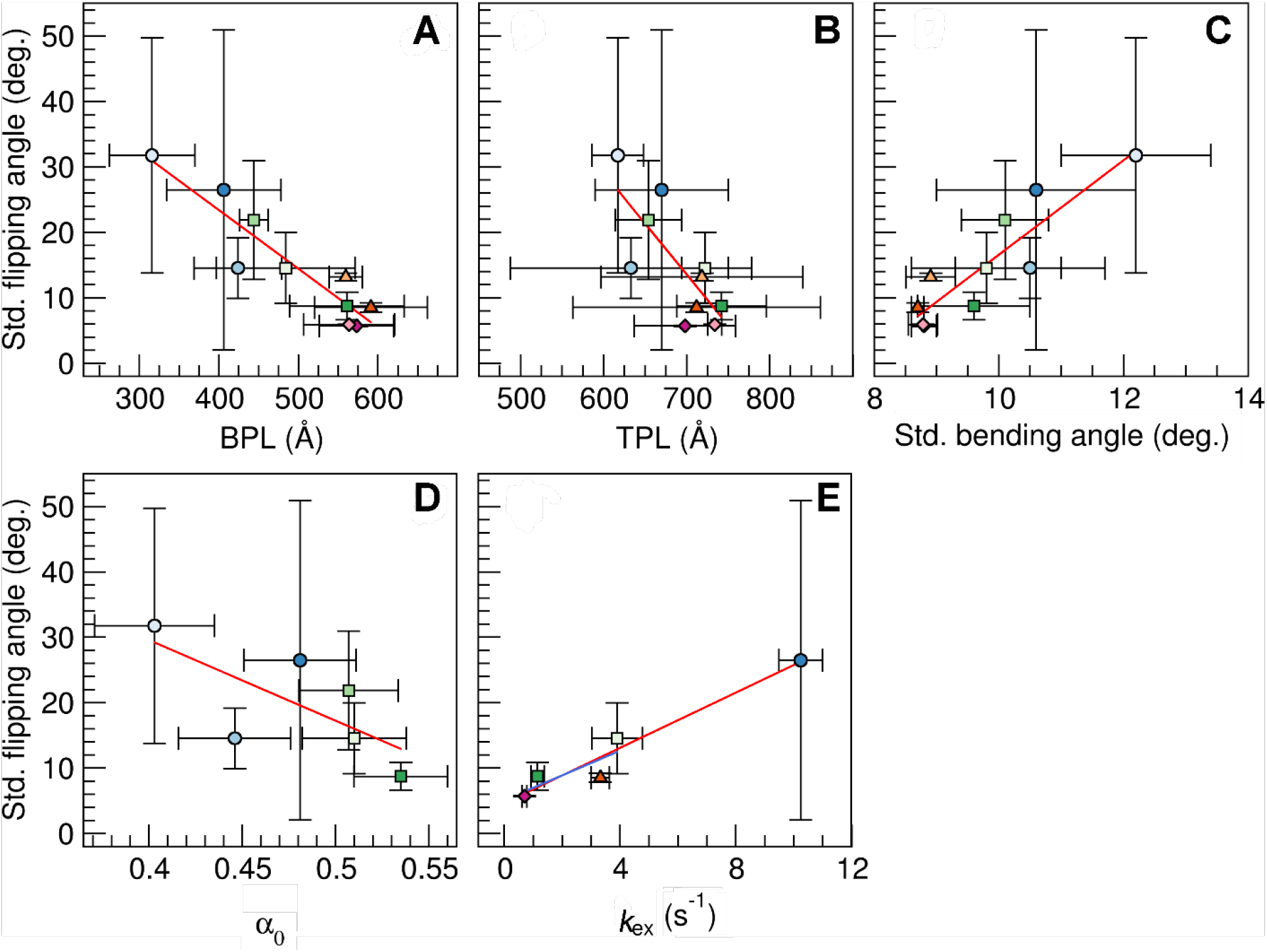
Correlation of the standard deviation of the flipping angle with the bending (A) and torsional persistence length (B), the standard deviation of the bending angle (C), α_0_ (D) and *k*_ex_ of U_6_ (E). Averages and standard deviations over 3 MD replicas. Red lines show linear regression of all data with correlation coefficients of −0.972 (A), −0.763 (B), 0.897 (C), −0.694 (D), and 0.972 (E), blue line the regression excluding 4TA with a correlation coefficient of 0.799 (E). 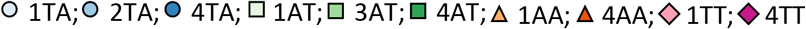.

### NMR-detected Imino Proton Exchange Rates

We probed individual base pair imino proton exchange rates of UNG substrates with solution NMR spectroscopy. Imino proton exchange rates are directly related to nucleic acid breathing motions, whereby the imino protons of each base pair in its transiently open state can exchange with water protons in the surrounding buffer (Fig. 5A) (42,70,71). The imino proton exchange rate (*k_ex_*) provides insight into the base pair stability and duplex dynamics at the single base-pair level. As indicated by the MD simulations, we hypothesized that the *k_ex_* of the target uracil (U_6_, Fig. 5B) will be sequence-dependent, and that a higher *k_ex_* for central base pairs reflects more suitable substrates for UNG. The imino protons for each NMR duplex listed in Table 1 were assigned (Fig. S3). The individual base pair *k_ex_* was then measured for each UNG substrate. The 1TA exchange rates were not characterized due to significant resonance overlap of uracil (U_6_) with T_18_ (Fig. S3) which impeded accurate deconvolution and analysis of the U_6_ and T_18_ resonances in *R*_1n_ and *k_ex_* experiments. The UNG substrates from series one were therefore not included in future NMR analyses.

**Figure 5.**
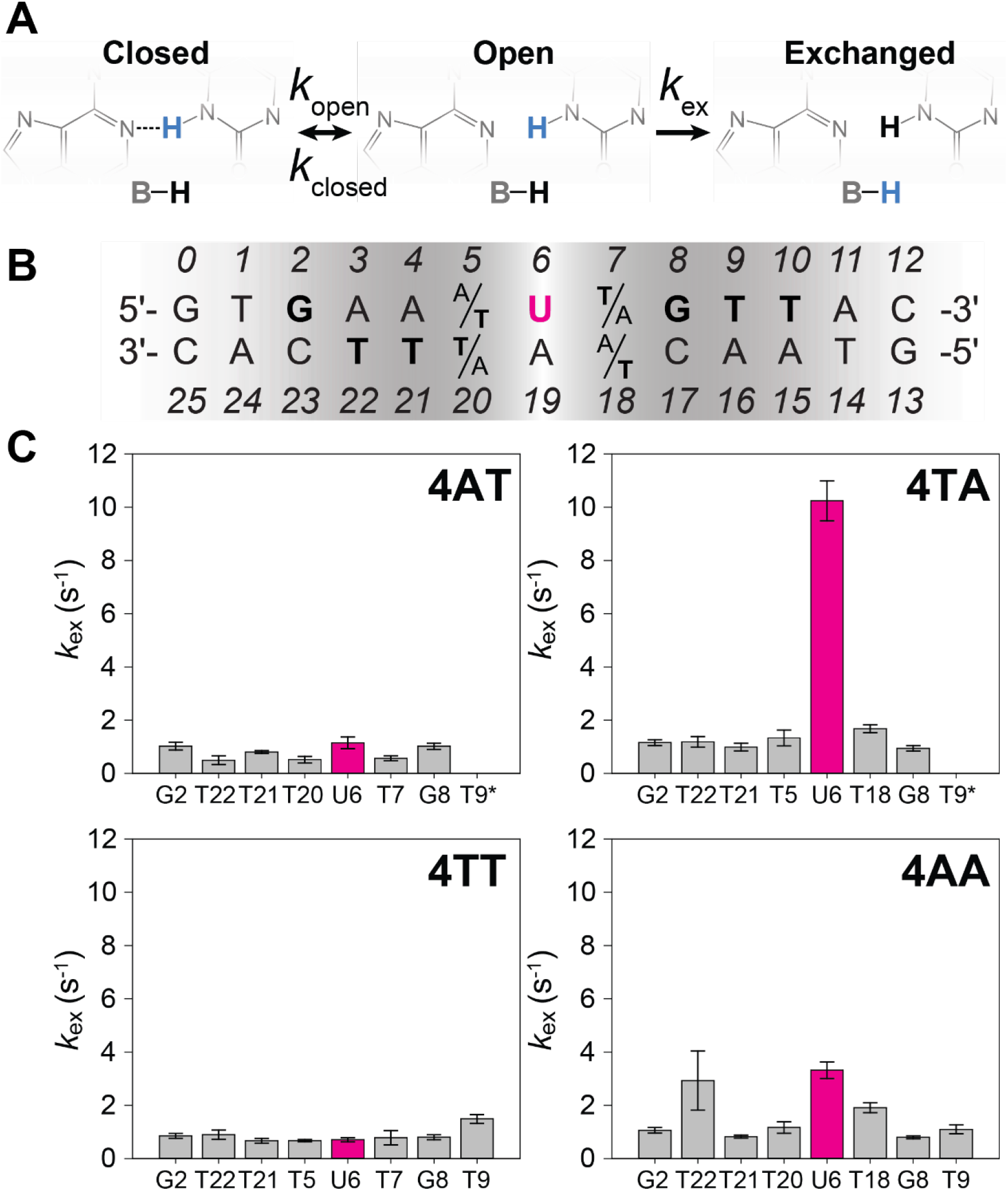
Imino proton exchange rates (*k*_ex_) of individual base pairs in each of the four-series substrates in isolation confirm MD and fluorescence dynamics studies. A) Schematic representation of the two-state model for base pair opening and imino exchange kinetics in nucleic acids. When the *k*_ex_ is not the rate-limiting step, the *k*_ex_ is directly proportional to the stability of the base pair and provides insight into DNA duplex dynamics. B) Series four UNG substrate sequences, with the central uracil (U_6_) highlighted in magenta. C) Imino proton exchange rates (*k*_ex_) of individual base pairs in 4AT, 4TA, 4TT, and 4AA substrates. The uracil exchange rate corresponds to substrate efficiency. Error bars represent the propagated fitting error.

Imino proton NMR spectra are sensitive to structure and dynamics and are only observed in the presence of base-pairing (72–74). Terminal duplex base-pairs were not observed, as expected, due to duplex fraying and their rapid exchange with buffer protons. Though most of the surrounding exchange rates were comparable between sequences, the central uracil of each sequence exhibited strikingly distinct *k_ex_* rates that varied by more than ten-fold in a sequence-dependent manner (Fig. 5C). Differences in imino exchange rates at the uracil position demonstrate a strong dependence on adjacent base pairs. In 4AT and 4TA, the *k_ex_* significantly increases as the result of a conserved change in sequence order of the surrounding base pairs. Given the significant differences in dynamics observed by fluorescence, MD simulations, and NMR experiments when swapping bases flanking the central uracil between AT and TA sequences identified, we measured imino proton exchange rates for 4TT and 4AA to elucidate whether the 5’ T or 3’ A relative to uracil dictates substrate flexibility and UNG suitability. Interestingly, the sequences with the highest uracil imino exchange rates are those with adenine 3’ to U_6_ (A_7_). The 4TA duplex measures the highest U_6_ *k_ex_* of 10.24 ± 0.75 s^−1^ while 4AA is intermediate with a *k_ex_* of 3.32 ± 0.31 s^−1^ (Fig 5C). Our experiments identify that the base pair at the 3’ side of uracil is most influential on the *k_ex_* of U_6_, demonstrated by the nearly 15-fold difference in *k_ex_* between 4TA and 4TT (10.24 ± 0.75 s^−1^ and 0.70 ± 0.08 s^−1^, respectively). This could suggest a structural and/or energetic hindrance to UNG efficiency in repairing such sequences (75,76). A positive correlation between *k_ex_* of U_6_ and *k*_cat_/*K*_M_ was observed (Fig. 2B), indicating that more flexible sequences have higher repair efficiencies.

### Thermodynamic cycle analysis

Given that the base identity 3’ to uracil most significantly contributes to substrate flexibility, we sought to identify if there was an allosteric coupling between 5’ and 3’ flanking positions to uracil. Thermodynamic cycle analysis of substrate series four identifies that there is coupling between these positions with a ΔG_coup_ of 3.9 ± 0.4 kJ/mol (Eq. 2 and Fig. 6). If our observation that substrate dynamics govern UNG activity is correct, then one should see coupling between the 5’ and 3’ flanking uracil positions in enzyme assays. To validate this, an analogous thermodynamic cycle with substrate series four was carried out using Δ*G*^‡^ values calculated from *k*_cat_/*K*_m_ measurements. The thermodynamic cycle based on enzyme kinetics measurements identified a coupling energy of 1.8 ± 0.6 kJ between uracil flanking positions. Thermodynamic theory dictates that this allosteric coupling should be preserved independent of the surrounding sequence. To evaluate this possibility, a thermodynamic cycle was carried out with substrate series one to evaluate the uracil flanking positions. The results of these studies identify the same coupling energetics as seen in the substrate series four, as expected. The outcomes of the thermodynamic cycle analysis by NMR and UNG enzymology are shown in Fig. 6.

**Figure 6:**
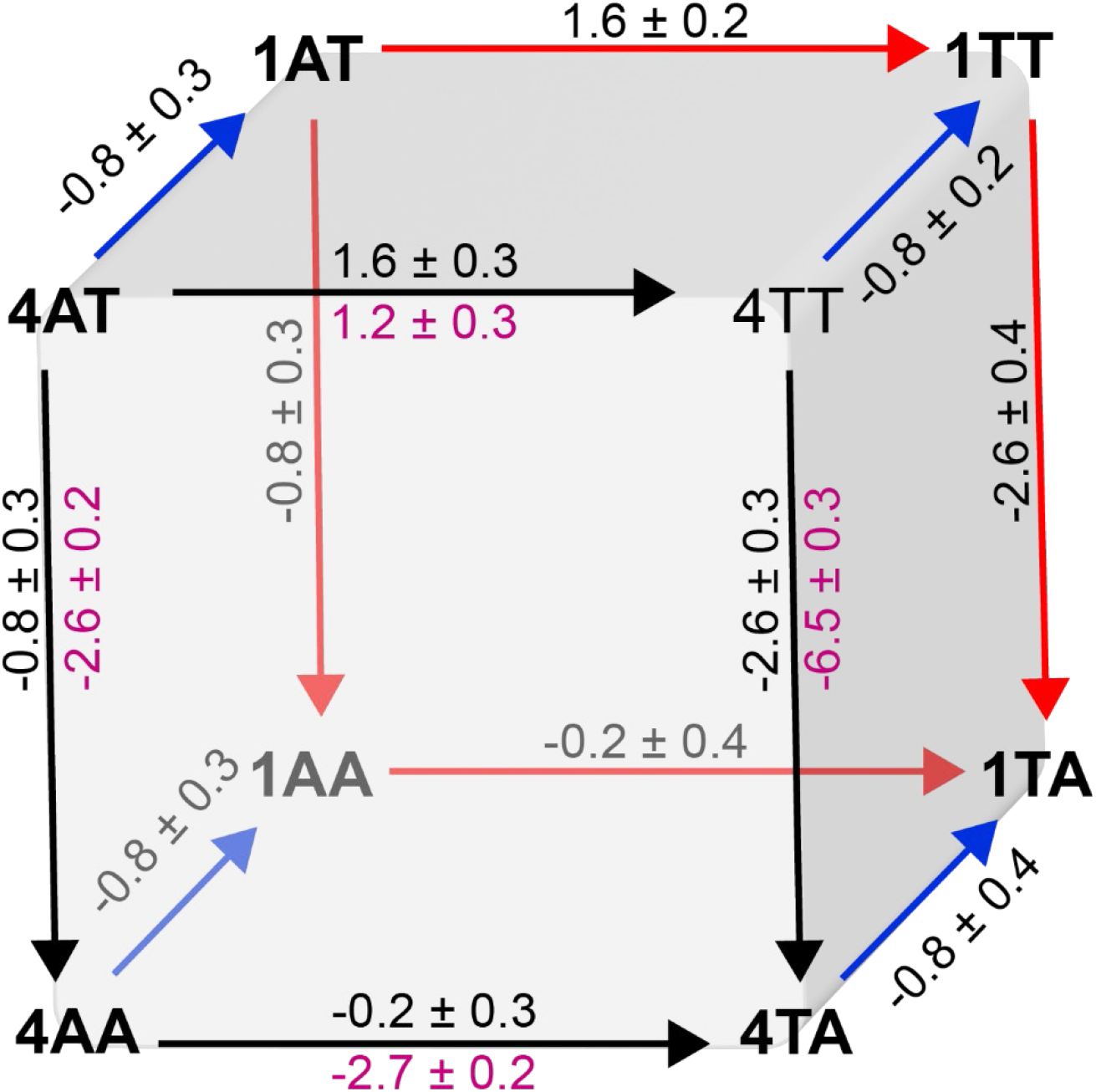
Double mutant cycle analysis for substrate series one (back face, red arrows) and substrate series four (front face, black arrows). All values represent 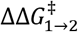 in kJ. Black values for substrate series one and four were calculated from *k*_cat_/*K*_m_ values of Table S2, magenta values for substrate series four from *k*_ex_ values of U_6_ as reported Fig. 5.

## DISCUSSION

Life on Earth depends on efficient duplication and subsequent transfer of DNA-based genetic material. Given that DNA is a dynamic and substantial biopolymer (each human cell contains ~3 billion nucleotides), the quality control and maintenance of DNA are central to biological fitness. Genomic integrity is maintained by diverse cellular mechanisms, including the Base Excision Repair (BER) pathway. BER is initiated by lesion recognition and excision by glycosylase enzymes that create an abasic site that is then processed in subsequent steps to preserve the DNA integrity. The most common human base substitution mutation is a C → T transition, which can arise from uracil in DNA (9,77). Uracil lesions are common in humans and across biology. The first uracil DNA glycosylase (UNG) was identified almost 50 years ago in *E. coli* (78), and since, the equivalent enzyme was identified across mammals, plants, bacteria, and some viruses. While UNG is well characterized functionally and structurally, a documented phenomenon where DNA substrate sequence impacts UNG functional efficiency, by orders of magnitude, has not been mechanistically explained. A better understanding of the fundamental principles that dictate UNG activity has broad implications in diverse fields, from cancer (79) to evolution (80). Our study employed a small library of DNA UNG substrate duplexes that were designed with variable efficiencies, enabling the deconvolution of the physical DNA substrate properties that impact UNG function. UNG activity was correlated to substrate uracil dynamics by fluorescence, NMR, and molecular dynamics investigations. These studies identified both proximal and distal substrate features whose contributions impact UNG activity.

UNG catalyzes uracil excision from duplex DNA by a base flipping mechanism in which the uracil rotates from the DNA base stack into the enzyme active site (18). NMR dynamics studies have established that uracil recognition by human and *E. coli* UNG relies on trapping extrahelical uracils that are spontaneously exposed by thermally induced base pair opening motions (81,82). The sequence context surrounding the uracil has been shown to affect the rate of uracil removal, but studies to date involved kinetic measurements on random substrate libraries that are not sufficient to identify the molecular origins of the observed differences (5,6,20,21). We have determined the UNG Michaelis-Menten constants for ten designed DNA substrates containing uracil flanked by either A or T bases (Fig. 1, Fig. S7, Table S2). This experimental design allowed us to focus on the effects of substrate flexibility in the vicinity of the uracil without affecting the melting temperature and stability of the duplex. UNG specificity constants (*k*_cat_/*K*_m_) were calculated and used to evaluate relative UNG substrate preferences. Relative *k*_cat_/*K*_m values_ were additionally determined from competition assays, and were found to agree with the relative efficiencies determined from the Michaelis-Menten parameters measured for substrates in isolation. We identify that the UNG specificity constant (*k*_cat_/*K*_m_) is generally smaller for substrates containing a thymine 3’ to the uracil. Replacing the 3’ thymine in substrates 1TT and 4TT with an adenine (TUT → TUA), results in a *c.a*. 3-fold increase in *k*_cat_/*K*_m_. Swapping A and T around the uracil in substrates 1AT and 4AT (AUT → TUA) also results in an increase in the specificity constant.

The fact that UNG recognizes spontaneously exposed uracil suggests that sequence effects in *k*_cat_/*K*_m_ are due to inherent differences in the deformability and flexibility of the DNA helix around the uracil. Accordingly, we observe an inverse correlation between *k*_cat_/*K*_m_ and α_0_, a fluorescence-derived experimental observable that measures the degree of base stacking in the region surrounding the uracil (Fig. 2A). Values of α_0_ are smaller for TUA sequences than for AUT, indicating that uracil is less dynamic in the second group. Similarly, a thymine 3’ to U results in lower exchange rate constants for the uracil imino proton (*k*_ex_, Table S1 and Fig. 2B), and we observed a correlation between *k*_cat_/*K*_m_ and *k*_ex_ for the five substrates for which we were able to obtain *k*_ex_ constants by NMR (Fig. 2B). This result further supports the notion that the UNG specificity constant is greater for substrates containing the uracil in more flexible contexts. We note that there is no clear correlation between *k*_cat_ and α_0_ (Fig. S9), indicating that the values of *K*_m_, but not *k*_cat_, are dictated by the mechanical properties of the substrate. This is consistent with current UNG mechanistic knowledge where the UNG *k_cat_* is limited by product release (83–85), so correlations with the properties of the uracil-containing substrate are not expected.

Our MD data suggests that the α_0_ values derived from time-resolved fluorescence spectroscopy and the U_6_-*k*_ex_ values determined by NMR measure different aspects of substrate motion. Whereas the correlation between α_0_ and bending motions was high (*r* of ~0.9), this correlation was less for *k*_ex_ of U_6_ (Fig. S10). When considering all sequences, the correlation coefficients with the bending persistence length, the torsional persistence length, and the standard deviation of the bending angle were −0.889, 0.739, and 0.788, respectively; however, since the *k*_ex_ values cluster, these large *r* values were mostly due to the outlying 4TA sequence. Without 4TA, *r* values of −0.504, 0.054, and 0.239 were obtained. At the same time, *k*_ex_ shows higher correlation with the standard deviation of the flipping angle (*r* of 0.972 for all sequences, and 0.799 when excluding 4TA) than α_0_ (*r* of −0.694). These data indicate that α_0_ values correlate more strongly with DNA bending than uracil flipping (Fig. 3, 4), while U_6_-*k*_ex_ values correlate more strongly to base flipping than DNA bending (Fig. 4, S10). Nevertheless, these motions are coupled (Fig. 4), and given the correlation of α_0_ and U_6_-*k*_ex_ with *k*_cat_/*K*_m_ (Fig. 2), both motions contribute to UNG substrate recognition and eventual uracil excision. The necessity of both motions for repair is consistent with human UNG structures, in which DNA substrate in complex with UNG has both a flipped uracil and bent duplex (by an average of 33°) (86–92). The stronger correlation of α_0_ with *k*_cat_/*K*_m_ (Fig. 2) is intriguing since it would indicate that excision is more strongly correlated to DNA bending than base flipping. The observation that *K*_m_, but not *k*_cat_, is governed by the mechanical properties of the substrate, indicates that DNA bending favors the formation of the enzyme-substrate complex, and the easier DNA is bent, the easier the enzyme-substrate complex forms. Our MD data shows that DNA bending correlates with base flipping, which is consistent with previous studies that show that DNA bending facilitates base flipping by pushing the system up in energy (93,94).

For a given substrate, changing one of the bases adjacent to uracil affects *k*_cat_/*K_m_* in a way that depends on the identity of the other one. For example, for substrate series one (i.e., 1AT, 1AA, 1TT, and 1TA) and four (i.e., 4AT, 4AA, 4TT, and 4TA), the effect of replacing an adenine 3’ to uracil with thymine is much greater when the base 5’ to uracil is T than A (e.g., compare 4AA→ 4AT vs. 4TA→ 4TT). Similarly, the effect of substituting an adenine 5’ to uracil by thymine is much greater when the base 3’ to uracil is T than A. The thermodynamic coupling between the bases flanking the uracil was analyzed by two types of double mutant cycles, one using data from solution NMR on the isolated substrates, and one from UNG activity (Fig. 6) (43). The first of these used the transition state free energies (Δ*G*^‡^) calculated from imino proton exchange rates for substrate series four (Fig. 6). Building on the identified substrate allosteric coupling, and our data linking function with substrate dynamics, we predicted that an analogous second thermodynamic cycle, this time using enzyme kinetic data, should parallel the cycle calculated from substrate dynamics. This prediction was validated by the second thermodynamic cycle, which used Δ*G*^‡^ values calculated from *k*_cat_/*K*_m_ ratios for substrate series one and four as shown in Fig. 6 (43,95). Since the cycles correspond to fundamentally different processes, the free energy values are expected to differ. Nevertheless, it is evident from both cycles that 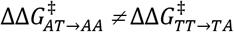 and 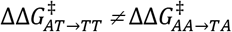, indicating that substituting one of the bases adjacent to uracil affects the energy of the transition state in a manner that depends on the identity of the other one. In other words, the sites directly 5’ and 3’ of uracil are allosterically coupled. Substituting a thymine 3’ to uracil by adenine (vertical edges of the cube in Fig. 6) stabilizes the transition states, but in both cycles, effects are more significant when the base 5’ to uracil is thymine. Similarly, substituting a thymine 5’ to uracil by adenine stabilizes the transition state when the base 3’ to uracil is thymine, but destabilizes it when adenine is in this position. This allosteric coupling is quantified by the coupling energy (Eq. 2), with a value of 3.9 ± 0.4 kJ/mol for substrate series four of the imino proton exchange-based cycle, and values of 1.8 ± 0.6 kJ/mol for the same series of substrates and 1.8 ± 0.5 kJ/mol for substrate series one of the *k*_cat_/*K*_m_ -based cycle. While magnitudes of coupling necessarily differ between the two cycles, the coupling has the same sign in both cases, which supports the conclusion that UNG catalysis correlates with substrate dynamics. Coupling energies are higher for the imino proton exchange-based cycle compared to the *k*_cat_/*K*_m_ -based cycle. Considering that the former measures coupling in the isolated substrates, these results point to the role of the enzyme in reducing the coupling energy, consistent with the fact that UNG must be capable of removing uracil in any sequence context.

The observation that the *k*_cat_/*K*_m_-based cycle coupling energies for substrate series one and four are identical, within error, confirms that positions proximal to the uracil lesion are fundamental in UNG efficiency. Furthermore, according to this cycle, the effect of a single change in a base adjacent to uracil is the same for substrates series one and four (i.e, 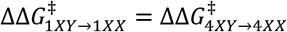, where X and Y are A or T).

This indicates that all the combined differences between substrates one and four have a constant effect in *k*_cat_/*K*_m_ that is independent of the identity of the bases flanking the uracil. Consistent with this, all values of 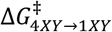 are the same for any combination of X and Y (lines connecting the thermodynamic squares of substrates 1 and 4 in Fig. 6). Although results show that sequence effects are not limited to the bases immediately surrounding the uracil lesion, for these sequences, effects appear to be additive to the effects of the bases flanking the uracil. For pairs of substrates series one and four containing the same flanking bases (i.e. 1AA vs 4AA, 1TA vs 4TA, etc), *k*_cat_/*K*_m_ values are 1.4-fold greater for substrate series one, consistent with a stabilization of the transition state of 0.8 ± 0.9 kJ/mol. We note that the one and four series substrates were designed from previous studies that reported them to be among the best and worst substrates in large and complex DNA contexts (6). As such, we conclude that uracil proximal sites have the most significant impact in both substrate flexibility and UNG activity. This flexibility manifests in UNG activity, essentially modulating enzyme efficiency.

The sequence of steps that lead to the formation of the catalytically-active enzyme-DNA complex has been described as a “pinch-pull-push” mechanism that starts with the distortion of the DNA backbone imposed by a “pinching” action of a trio of serine residues (96, 97). Uracil is then displaced from the duplex (“pull”) and stabilized in an extrahelical, flipped-out conformation by the insertion of Leu191 into the formed DNA cavity (“push”). Ultimately this facilitates N-glycosylic bond cleavage to produce an abasic (AP) site and a free uracil base. The “push” step, insertion of Leu191, acts as a ‘doorstop’ that prevents the return of the flipped-out uracil residue. The first two steps, and possibly the third, require conformational changes in the DNA substrates, including hydrogen bond breakage and loss of stabilizing base stacking interactions. Considering that these interactions are determined by the sequence-dependent mechanical properties of the DNA, we hypothesize, supported by our fluorescence, NMR and MD data, that substrate flexibility defines the energetic barriers of the kinetic steps that precede cleavage. Future transient-state kinetic experiments that are beyond the scope of this work could dissect the effect of substrate flexibility on each of these individual steps.

While our study focused on the best-studied member of the UNG family, *E. coli* UNG, we anticipate the results to be broadly applicable because the catalytic cores of UNGs from different sources are closely related. For example, the root-mean-square deviation between all Cα positions of human and *E. coli* UNG enzymes is just 0.9 Å (98,99). These enzymes share an overall sequence identity of 56%, which includes conserved catalytic water activating loops (62-GQDPYH-67), the uracil specificity region (120-LLLN-123), and active site residues (D64, Y66, F77, N123, H187, and L191) (100). Structural similarities among uracil-DNA glycosylases transcend the UNG family. UNGs are structurally similar (despite low sequence identity) to bacterial mismatch-specific uracil-DNA glycosylases (MUGs) and to eukaryotic thymine-DNA glycosylates (TDGs). These enzymes recognize G:U/T lesions (100,101), and as UNGs, show a base-flipping mechanism for the recognition of uracil and thymine. We speculate that the MUG and TDG efficiencies will be dictated in large part by substrate lesion DNA dynamics, though future studies are needed to test this hypothesis.

In eukaryotes, most repair occurs on DNA that is packaged by histones into nucleosomes. While nucleosomes inhibit the activity of some glycosylases (e.g. hOGG1 and Fpg) (102), UNG removes uracil from solvent-exposed locations with an efficiency approaching that of naked DNA (102–105). These locations include the DNA outside the nucleosome core region, and sites in the nucleosome with outward rotational positioning. Repair of uracils oriented toward the histone core occurs at a slower rate, and relies on the formation of transient states that locally release the DNA (102–106). Considering that UNG removes uracil from either solvent-exposed locations or transiently exposed states, we speculate that the sequence effects we have observed with naked DNA substrates apply to packaged DNA.

Given the fundamental nature of genomic integrity, the implications of our studies are significant. That repair efficiencies, at least for UNG, are in large part dictated by DNA sequence deformability and flexibility could help explain the molecular mechanisms that underly fundamental observations in diverse fields. These include oncogenetics and cancer hotspots (79), and evolutionary adaptation (80), among other fields. One particularly relevant context extends to the rapidly expanding field of base editing, where UNG and other glycosylases have been tethered to Cas9 nickases enabling precision DNA alterations with great potential for therapeutic intervention (105,106). For instance, a new class of editors consisting of a Cas9 nickase, UNG, and a cytidine deaminase was shown to enable C-to-G base transversions (108). Uracils formed by deamination of a cytosine are removed by UNG, creating an abasic site that is then filled preferentially by a G. The mechanism behind this preference is still unknown, but C-to-G transversion was shown to be sequence dependent, and most efficient for Cs embedded in an AT-rich sequence. This finding is intriguing in light of our observation that UNG activity correlates with substrate flexibility. However, establishing whether UNG sequence preferences ultimately dictate C-to-G editing efficiency or not would require a deeper mechanistic understanding of how C-to-G edits are induced in these systems. Ultimately, our data show a clear correlation between UNG activity and substrate flexibility that can be used to make predictions about the functional attributes of substrates, and may help rationalize sequence effects in this and other fields.

## SUPPLEMENTARY DATA

Supplementary Data are available at NAR Online.

## ACKNOWLEDGEMENT

M.L. acknowledges use of the Ultrafast Laser Spectroscopy Facility at Arizona State University. W.V.H. acknowledges use of the Magnetic Resonance Research Center at Arizona State University. AvdV acknowledges use of the USF Research Computing facility.

