## Supplemental materials for "Uracil-DNA glycosylase efficiency is modulated by substrate rigidity"

**Supplemental Information: Uracil-DNA glycosylase efficiency is modulated by substrate rigidity**

**Table of Contents**


| | Page |
|---|---|
| Structure of 2-Aminopurine incorporated into oligonucleotides. | S2 |
| Sample kinetic run and a $V_0$ calculation and Fig. S1 | S2 |
| Figure S2 | S3 |
| Figure S3 | S3 |
| Water inversion efficiency factor (E) and Fig. S4 | S4 |
| Longitudinal relaxation of water ($R_{1w}$) and Fig. S5 | S4-S5 |
| Imino proton longitudinal relaxation ($R_{1n}$) and exchange rates ($k_{ex}$) | S6 |
| Figure S6 | S7 |
| Table S1 (Fitting parameters for imino proton exchange rate measurement ($k_{ex}$)) | S7-S8 |
| Figure S7 | S9 |
| Table S2 (Michaelis-Menten parameters ($K_m$ and $V_{max}$)) | S10 |
| Table S3 (Fluorescence quantum yields) | S11 |
| Table S4 (Fluorescence lifetimes) | S11 |
| Table S5 ($\alpha_0$ values) | S11 |
| Table S6 (MD properties) | S12 |
| Figure S8 (Step parameters for central $X_5U_6Y_7$ steps) | S12 |
| Figure S9 | S13 |
| Figure S10 (Correlation of MD properties with $k_{ex}$.) | S13 |

**2AP substitution** Structure of 2-Aminopurine incorporated into oligonucleotides.

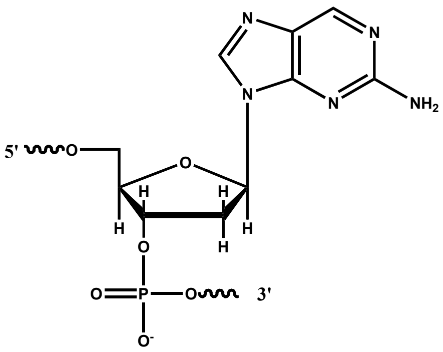

**Sample kinetic run and a $V_0$ calculation**

**Figure S1.** Fluorescence intensity (arbitrary units) vs time before and after the addition of UDG. The figure shows the results of a kinetic run using 1μM **1TA** DNA in 1x PBS buffer and 0.16 nM UDG. The measurement was interrupted briefly to add UDG and mix.

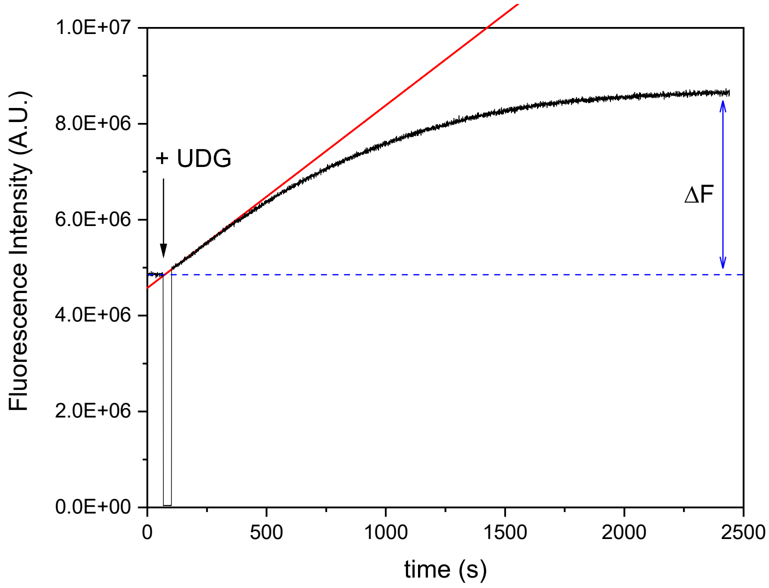


The initial velocity is calculated according to

$$V_0 = \frac{[S]_0}{\Delta F}\left(\frac{dF}{dt}\right)_0$$

Where $[S]_0 = 1.0 \times 10^{-6}$ M is the concentration of substrate, $\Delta F = 3.80 \times 10^6$ A.U is the difference between the final ($t \rightarrow \infty$) and initial Fluorescence intensity, and $\left(\frac{dF}{dt}\right)_0 = 3810\ A.U/s$ is the initial slope of the *F*(*t*) vs *t* graph. A.U. denotes arbitrary units. A value of $V_0 = 1.0 \times 10^{-9} \mathrm{M}.s^{-1}$ was calculated using data measured in this experiment.

**Figure S2** Two-dimensional $^{1}H$, $^{1}H$ - Nuclear Overhauser Effect Spectroscopy (NOESY) experiments were measured at 20 °C. Resulting 2D spectra were used to assign imino protons using traditional "backbone-walking" methods.[1,2]

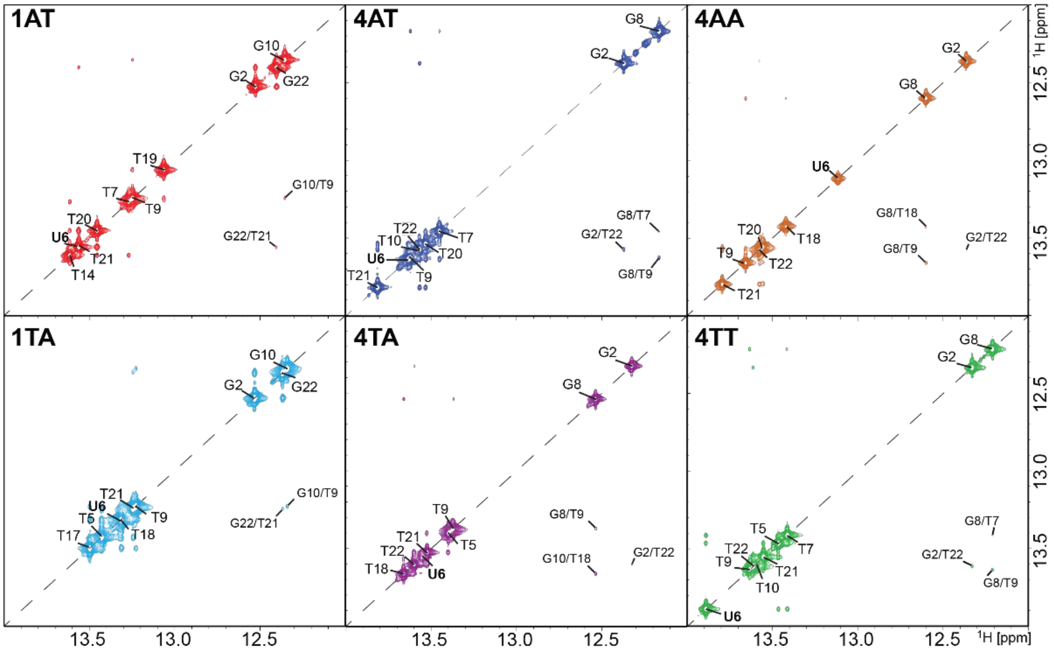


**Figure S3** A $^{1}H$ one-dimensional representation of the imino assignments determined from NOESY spectra in Fig. S2. Accompanying sequences include assigned imino bases in **bold**. All $^{1}H$-1D experiments were measured at 20°C.

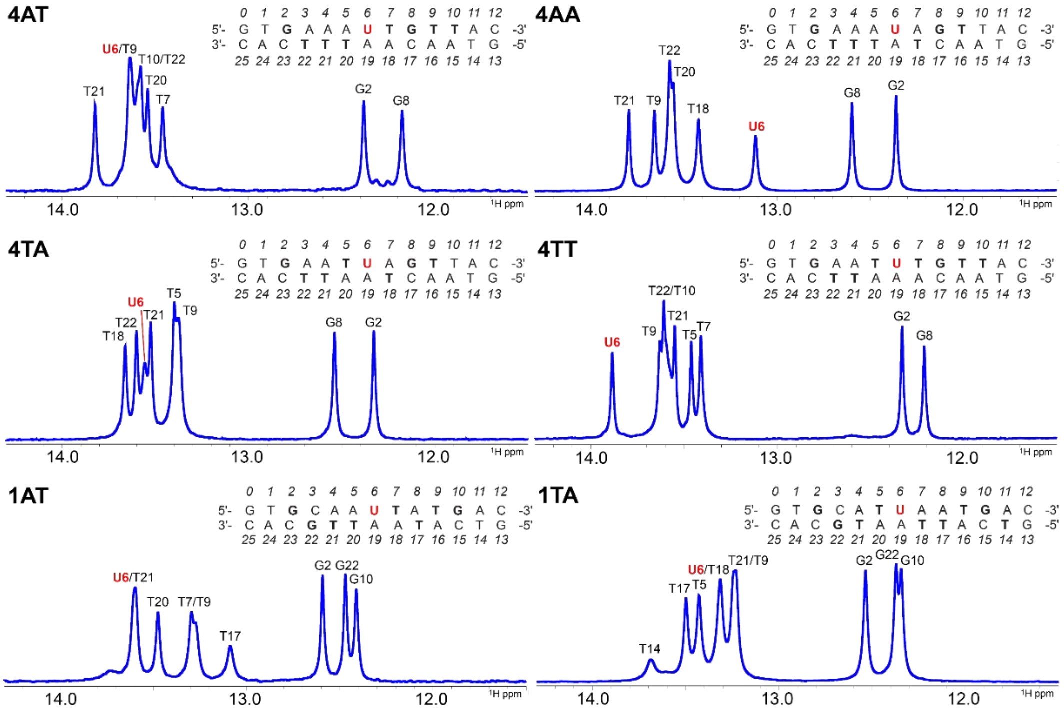

### Water inversion efficiency factor (E)

The water inversion efficiency factor *(E)* was determined in a 3 mm NMR tube on both the 600 MHz (Fig. S4A) and 850 MHz (Fig. S42B) instruments. The peak areas for each water signal were determined by manually phasing the spectra in Topspin 4.1 and MestReNova 14.2, respectively, and measuring the peak integrals in an overlayed spectrum. The measured peak areas were applied to the following equation

$$E = 1 - \frac{W_{inv}}{W_{eq}} \qquad \text{Eq. S1}$$

where $W_{inv}$ is the peak area of the signal after inversion and $W_{eq}$ is the peak area of the signal without inversion. The resulting *E* factor represents the efficiency of the selective pulse inversion, where *E* = 2 indicates total inversion efficiency. Previously reported *E* factors range from 1.90-2.00.[3] However, since the efficiency is mostly dependent on instrument hardware, these reference values could be inconstant. The measured *E* factor for the 600 MHz and 850 MHz were 1.905 and 1.943, respectively (Fig. S4C-D).

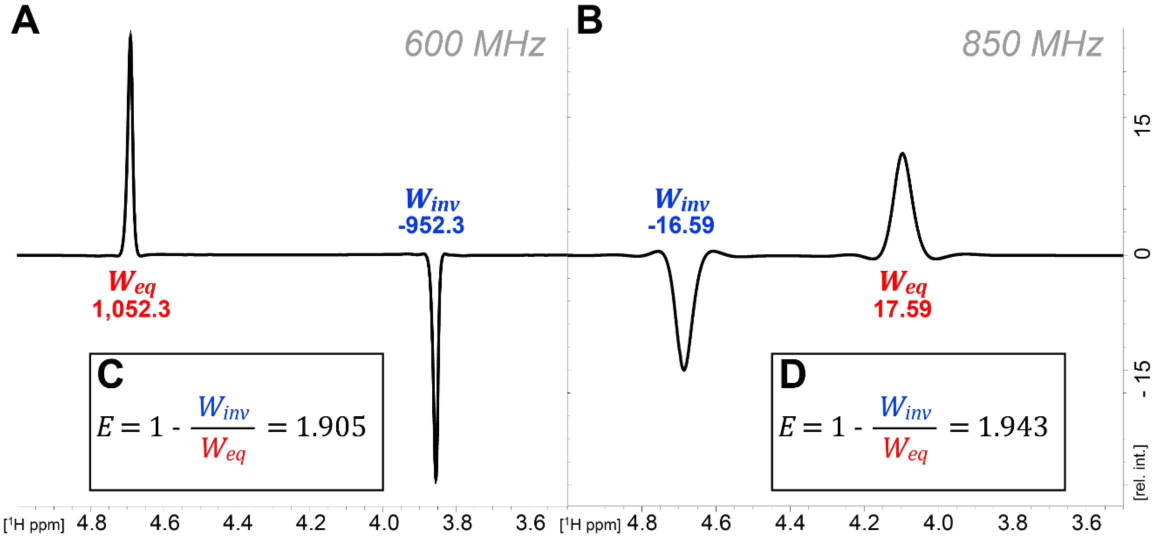


**Fig. S4**. Determination of the water inversion efficiency factor (*E*). A) The water signal peak area with ($W_{inv}$, blue) and without ($W_{eq}$, red) a DANTE inversion pulse[3,4] measured on the 600 MHz and B) the 850 MHz instruments. Spectra are overlayed and referencing is shifted 500 Hz for each. C) Applying Equation #, the measured *E* factor for the 600 MHz instrument is 1.905 and for D) the 850 MHz instrument it is 1.943.

### Longitudinal relaxation of water ($R_{1w}$)

The longitudinal relaxation of water ($R_{1w}$) was measured on the 600 MHz instrument in a 3 mm NMR tube using a list of 24 variable time delays ranging from 1 ms to 18 s and a relaxation delay of 30 s. Determination of $R_{1w}$ was initially completed using the TopSpin T1/T2 Module and fitting to the equation

$$A = \alpha + \beta e^{-tR_{1w}} \qquad \text{Eq. S2}$$

where *A* is the area of the water signal peak, *t* is the relaxation delay time, and α and β are constants. The TopSpin module resulted in a $R_{1w}$ of 0.5051. This value was verified in MATLAB R2019b (Mathworks) Curve

Fitting Tool to be 0.504 ± 0.012 (Fig. S5). Multiple trials were performed on the 600 MHz and 850 MHz magnets, which resulted in a working range of 0.4985 to 0.5102 for the $R_{1w}$ parameter in $k_{ex}$ fitting.

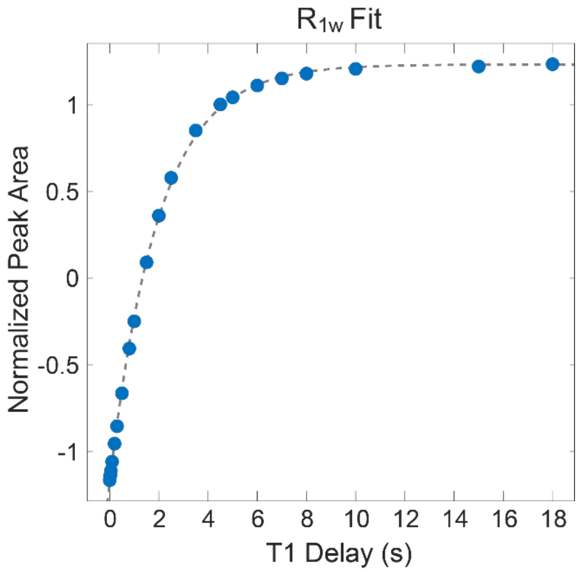


**Fig. S5.** Determination of the water longitudinal relaxation rate ($R_{1w}$). The water signal peak areas (blue) as a function of T1 delay was fit to Equation S2 (gray, dashed line) and rendered a $R_{1w}$ value of 0.5051.

**Imino proton longitudinal relaxation ($R_{1n}$) and exchange rates ($k_{ex}$)**

The longitudinal relaxation of water ($R_{1n}$) was initially measured on the 850 MHz instrument in a 3 mm NMR tube using a list of 24 variable time delays ranging from 1 ms to 15 s. However, imino proton peaks in some sequences exhibited low resolution and S/N despite the high-field instrument. For these sequences, data was collected on the 600 MHz instrument where resolution and S/N significantly improved, presumably because of more favorable relaxation properties. Previously described pseudo two-dimensional experiments[4] for the imino proton $R_{1n}$ and imino proton exchange rate ($k_{ex}$) were measured on the same instrument for each sequence.

The respective spectra were processed and phased in TopSpin 4.1. The data were baseline corrected and fit with MATLAB R2019b (MathWorks) using nonlinear least squares fit (Fig. S6A-D). The $R_{1n}$ was determined by fitting the individual imino proton peaks areas to the equation:

$$A = \alpha + \beta e^{-tR_{1n}} \quad \text{Eq. S3}$$

where $A$ is the area under the peak, $t$ is the relaxation delay time, and α and β are constants (Fig. S6E). The error of the fit was used as the variable bounds for the $R_{1n}$ parameter in fitting for $k_{ex}$. The ($k_{ex}$) was determined by fitting the individual imino proton peak areas to Equation 1 (main manuscript) (Fig. S6F). The $R_{1w}$ and $R_{1n}$ parameters for fitting $k_{ex}$ were permitted to float within the error bounds previously determined in the respective fits, whereas the $E$ parameter was fixed for data collected on the 600 MHz and 850 MHz, respectively (Table S1). The final $k_{ex}$ for each imino proton was reported with the error of the fit.

**Figure S6** Example of determining the imino proton longitudinal relaxation ($R_{1n}$) and exchange ($k_{ex}$) rates from 1AT. A) Spectra processed and phased in TopSpin 4.1 were baseline corrected in MATLAB R2019b (MathWorks). B) Peaks were fit to the new baseline with a Lorentzian function and C) individual peaks were fit based on predetermined peak positions. D) Resulting residuals of individual peak fitting. E) $R_{1n}$ fitting of the 1AT G2 imino proton peak areas to Equation S3. F) $k_{ex}$ fitting of the 1AT G2 imino proton peak areas to Equation 1 (main manuscript).

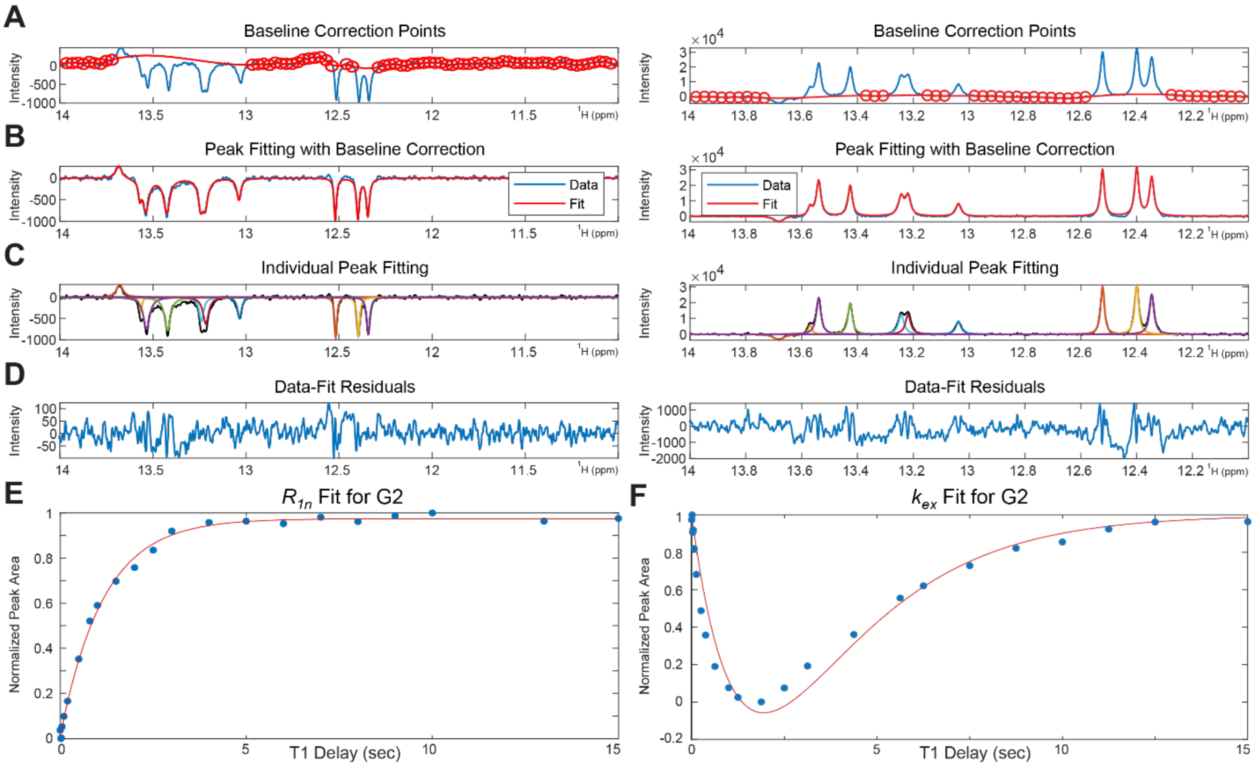


**Table S1.** Fitting parameters for imino proton exchange rate measurement ($k_{ex}$)

| 4AT | | | | | | |
|---|---|---|---|---|---|---|
| Imino | $k_{ex}$ ($s^{-1}$) | Error | $R_{1n}$ ($s^{-1}$) | Error | $^1$H ppm | σ |
| G2 | 1.03 | 0.15 | 0.819 | 0.102 | 12.3566 | 0.0001 |
| T22 | 0.50 | 0.16 | 0.294 | 0.281 | 13.5537 | 0.0045 |
| T21 | 0.81 | 0.06 | 0.626 | 0.077 | 13.7983 | 0.0001 |
| T20 | 0.52 | 0.12 | 0.252 | 0.122 | 13.5161 | 0.0001 |
| **U6** | 1.15 | 0.22 | 1.197 | 0.449 | 13.6164 | 0.0011 |
| T7 | 0.57 | 0.09 | 0.352 | 0.161 | 13.4360 | 0.0003 |
| G8 | 1.02 | 0.12 | 0.892 | 0.101 | 12.1508 | 0.0001 |
| T9 | 7.02 | 2.25 | 9.593 | 3.922 | 13.6046 | 0.0044 |
| 4TA | | | | | | |
| Imino | $k_{ex}$ ($s^{-1}$) | Error | $R_{1n}$ ($s^{-1}$) | Error | $^1$H ppm | σ |

| | | | | | | |
|---|---|---|---|---|---|---|
| G2 | 1.16 | 0.11 | 1.040 | 0.097 | 12.3079 | 0.0001 |
| T22 | 1.19 | 0.20 | 1.065 | 0.321 | 13.5813 | 0.0003 |
| T21 | 0.99 | 0.15 | 0.844 | 0.275 | 13.5051 | 0.0002 |
| T5 | 1.34 | 0.30 | 1.238 | 0.215 | 13.3776 | 0.0004 |
| **U6** | 10.24 | 0.75 | 14.923 | 1.838 | 13.5373 | 0.0067 |
| T18 | 1.68 | 0.15 | 1.827 | 0.298 | 13.6429 | 0.0003 |
| G8 | 0.94 | 0.10 | 0.774 | 0.050 | 12.5195 | 0.0002 |
| T9 | 1.60 | 0.19 | 1.646 | 0.262 | 13.3521 | 0.0013 |
| 4AA | | | | | | |
| Imino | $k_{ex}$ ($s^{-1}$) | Error | $R_{1n}$ ($s^{-1}$) | Error | $^1$H ppm | σ |
| G2 | 1.06 | 0.10 | 0.884 | 0.110 | 12.4062 | 0.0001 |
| T22 | 2.93 | 1.11 | 3.569 | 2.458 | 13.6130 | 0.0061 |
| T21 | 0.82 | 0.06 | 0.603 | 0.057 | 13.8232 | 0.0001 |
| T20 | 1.17 | 0.22 | 1.096 | 0.413 | 13.5661 | 0.0007 |
| **U6** | 3.32 | 0.31 | 4.285 | 0.991 | 13.1387 | 0.0002 |
| T18 | 1.91 | 0.18 | 2.109 | 0.755 | 13.4357 | 0.0022 |
| G8 | 0.79 | 0.06 | 0.580 | 0.020 | 12.6260 | 0.0001 |
| T9 | 1.09 | 0.17 | 1.009 | 0.177 | 13.6849 | 0.0003 |
| 4TT | | | | | | |
| Imino | $k_{ex}$ ($s^{-1}$) | Error | $R_{1n}$ ($s^{-1}$) | Error | $^1$H ppm | σ |
| G2 | 0.85 | 0.09 | 0.618 | 0.024 | 12.3291 | 0.0001 |
| T22 | 0.89 | 0.17 | 0.718 | 0.123 | 13.6090 | 0.0017 |
| T21 | 0.67 | 0.09 | 0.438 | 0.069 | 13.5494 | 0.0002 |
| T5 | 0.67 | 0.04 | 0.460 | 0.083 | 13.4608 | 0.0000 |
| **U6** | 0.70 | 0.08 | 0.475 | 0.133 | 13.8850 | 0.0001 |
| T7 | 0.78 | 0.27 | 0.569 | 0.044 | 13.4095 | 0.0005 |
| G8 | 0.80 | 0.09 | 0.564 | 0.026 | 12.2099 | 0.0001 |
| T9 | 1.49 | 0.17 | 1.593 | 0.363 | 13.6282 | 0.0058 |
| 1AT | | | | | | |
| Imino | $k_{ex}$ ($s^{-1}$) | Error | $R_{1n}$ ($s^{-1}$) | Error | $^1$H ppm | σ |
| G2 | 1.02 | 0.11 | 0.885 | 0.089 | 12.5217 | 0.0002 |
| G22 | 0.89 | 0.11 | 0.765 | 0.060 | 12.3989 | 0.0003 |
| T21 | 0.63 | 0.14 | 0.342 | 0.081 | 13.5386 | 0.0006 |
| T20 | 1.12 | 0.15 | 1.031 | 0.105 | 13.4259 | 0.0002 |
| **U6** | 3.90 | 0.88 | 5.071 | 0.218 | 13.5732 | 0.0070 |
| T7 | 1.32 | 0.12 | 1.246 | 0.247 | 13.2430 | 0.0003 |
| T17 | 1.56 | 0.15 | 1.655 | 0.326 | 13.0378 | 0.0003 |
| T9 | 1.08 | 0.07 | 1.017 | 0.102 | 13.2189 | 0.0003 |
| G10 | 1.29 | 0.13 | 1.235 | 0.124 | 12.3455 | 0.0002 |

**Figure S7.** Experimental initial rates and Michaelis-Menten plots for samples 2AT, 4TA, 1AT, 3AT, 1AA, 4AA, 1TT and 4TT. The corresponding graphs for samples 1TA and 4AT are shown in Fig. 1. Michalis-Menten parameters are shown in Table S2. UDG concentration was 0.16 nM in all cases.

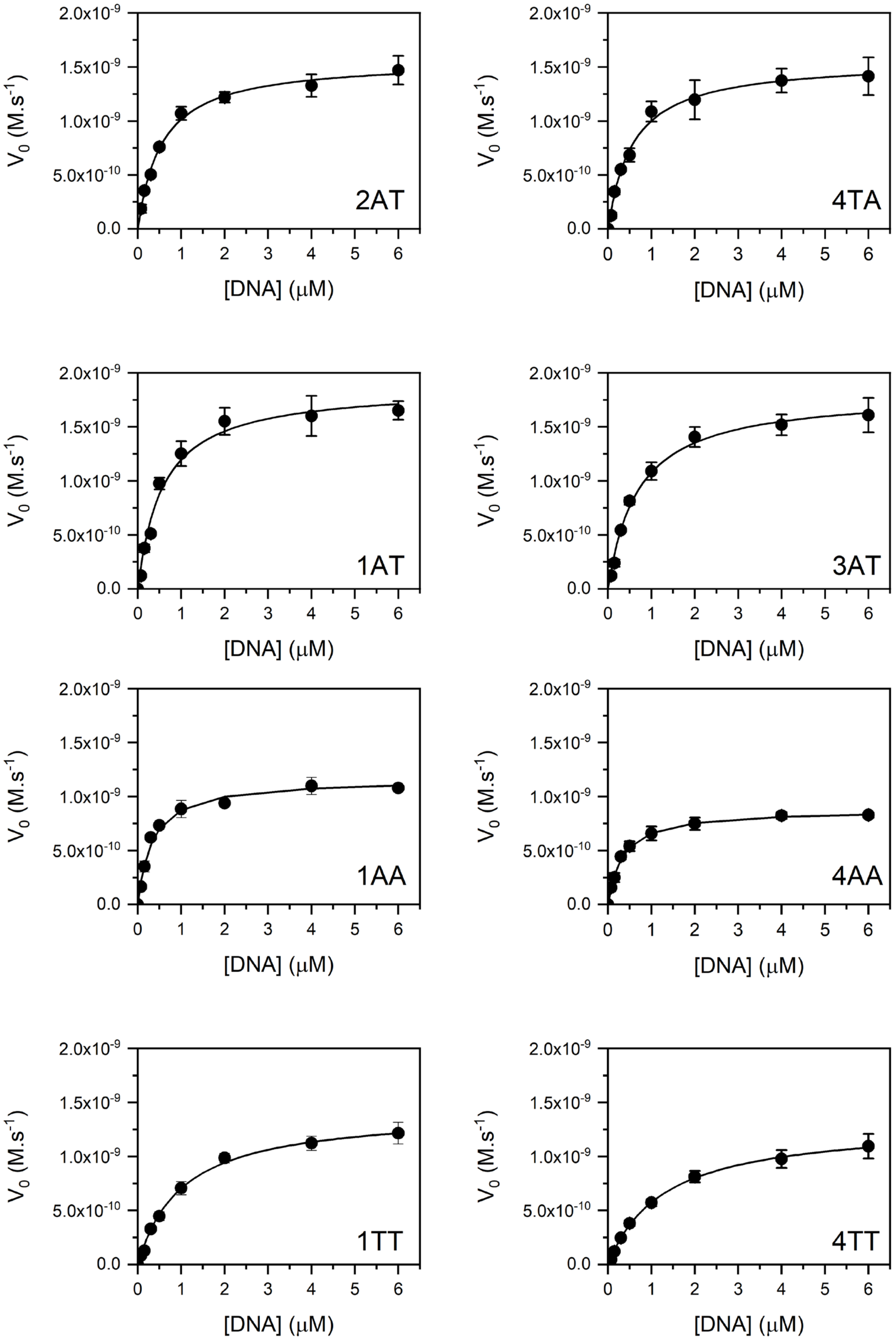

**Table S2** Michaelis-Menten parameters (Km and Vmax) where obtained by fitting the data shown in Figs. 1 and S7 to the following equation: $V_0 = \frac{V_{max}[S]}{K_m+[S]}$. The catalytic constant, $k_{cat}$, was calculated as $V_{max}/[E]$, where [E], the enzyme concentration, was calculated using the specific activity provided by the manufacturer (100 U/μg). Errors in Table S2 are calculated from the standard errors of the fitting parameters. Fitting was done using the Lavenberg Marquardt iteration algorithm using the reciprocal of the variances of each point as weights.

| Sample | $K_M$ (μM) | $k_{cat}$ ($s^{-1}$) | $\frac{k_{cat}}{K_M}$ ($M^{-1}$ $s^{-1}$) |
|---|---|---|---|
| 1TA | 0.356 ± 0.047 | 8.33 ± 0.25 | (2.34 ± 0.32) × $10^7$ |
| 1AT | 0.543 ± 0.024 | 8.731 ± 0.090 | (1.608 ± 0.073) × $10^7$ |
| 2TA | 0.524 ± 0.058 | 9.35 ± 0.20 | (1.78 ± 0.22) × $10^7$ |
| 3AT | 0.642 ± 0.059 | 10.7 ± 0.22 | (1.67 ± 0.15) × $10^7$ |
| 4TA | 0.538 ± 0.061 | 9.26 ± 0.23 | (1.72 ± 0.20) × $10^7$ |
| 4AT | 0.611 ± 0.058 | 7.09 ± 0.16 | (1.16 ± 0.11) × $10^7$ |
| 4AA | 0.332 ± 0.018 | 5.282 ± 0.072 | (1.589 ± 0.088) × $10^7$ |
| 4TT | 1.286 ± 0.061 | 7.89 ± 0.10 | (0.614 ± 0.030) × $10^7$ |
| 1AA | 0.321 ± 0.029 | 6.99 ± 0.15 | (2.18 ± 0.20) × $10^7$ |
| 1TT | 1.016 ± 0.057 | 8.53 ± 0.13 | (0.840 ± 0.049) × $10^7$ |

**Table S3** Fluorescence quantum yields (ϕ) were determined as described in Materials and Methods. Values are the average of at least 4 determinations. All standard deviations are 3% or less.

| Sample | 1TA | 1AT | 2TA | 3AT | 4TA | 4AT |
|---|---|---|---|---|---|---|
| ϕ | 0.00383 | 0.0103 | 0.00308 | 0.00981 | 0.00356 | 0.00606 |

**Table S4** Results of two independent TCSPC experiments. Time-resolved fluorescence intensity decays were acquired and fitted as discussed in Materials and Methods. $F(t) = \sum_{i=1}^{4} \alpha_i e^{-t/\tau_i}$, $\sum_{i=1}^{4} \alpha_i = 1$, $\langle\tau\rangle = \sum_{n=1}^{4} \alpha_i \tau_i$

Experiment 1

| Sample | $\alpha_1$ | $\tau_1$ (ns) | $\alpha_2$ | $\tau_2$ (ns) | $\alpha_3$ | $\tau_3$ (ns) | $\alpha_4$ | $\tau_4$ (ns) | $\langle\tau\rangle$ (ns) |
|---|---|---|---|---|---|---|---|---|---|
| 1TA | 0.709 | 0.0339 | 0.278 | 0.117 | 0.00937 | 1.46 | 0.00394 | 5.83 | 0.0932 |
| 1AT | 0.402 | 0.0662 | 0.495 | 0.312 | 0.0969 | 0.999 | 0.00624 | 6.09 | 0.316 |
| 2TA | 0.873 | 0.0326 | 0.116 | 0.144 | 0.00686 | 1.87 | 0.00408 | 5.92 | 0.0821 |
| 3AT | 0.381 | 0.0780 | 0.557 | 0.307 | 0.0556 | 1.05 | 0.00629 | 5.55 | 0.294 |
| 4TA | 0.676 | 0.0387 | 0.310 | 0.128 | 0.00970 | 1.66 | 0.00377 | 4.93 | 0.100 |
| 4AT | 0.593 | 0.0644 | 0.375 | 0.244 | 0.0253 | 1.082 | 0.00607 | 5.95 | 0.193 |

Experiment 2

| Sample | $\alpha_1$ | $\tau_1$ (ns) | $\alpha_2$ | $\tau_2$ (ns) | $\alpha_3$ | $\tau_3$ (ns) | $\alpha_4$ | $\tau_4$ (ns) | $\langle\tau\rangle$ (ns) |
|---|---|---|---|---|---|---|---|---|---|
| 1TA | 0.725 | 0.0366 | 0.263 | 0.123 | 0.00905 | 1.68 | 0.00365 | 6.98 | 0.0995 |
| 1AT | 0.420 | 0.0693 | 0.467 | 0.308 | 0.107 | 0.919 | 0.00642 | 6.68 | 0.315 |
| 2TA | 0.849 | 0.0328 | 0.140 | 0.122 | 0.00769 | 1.53 | 0.00405 | 6.59 | 0.0833 |
| 3AT | 0.384 | 0.0854 | 0.549 | 0.310 | 0.0603 | 0.982 | 0.00655 | 6.13 | 0.302 |
| 4TA | 0.675 | 0.0386 | 0.311 | 0.125 | 0.0106 | 1.69 | 0.00391 | 5.85 | 0.106 |
| 4AT | 0.574 | 0.0653 | 0.397 | 0.245 | 0.0230 | 1.20 | 0.00529 | 6.60 | 0.197 |

**Table S5** Fractional concentration of highly stacked 2AP ($\alpha_0$). Values of $\alpha_0$were calculated as $\alpha_0 = 1 - \frac{\tau_{2AP}}{\langle\tau\rangle}\frac{\phi}{\phi_{2AP}}$, using the values of Tables S3 and S4 and $\phi_{2AP} = 0.68$ and $\tau_{2AP} = 10.2$ ns (see Materials and Methods)

| Sample | 1TA | 1AT | 2TA | 3AT | 4TA | 4AT |
|---|---|---|---|---|---|---|
| $\alpha_0$ | 0.403 ± 0.032 | 0.510 ± 0.028 | 0.446 ± 0.030 | 0.507 ± 0.027 | 0.481 ± 0.030 | 0.535 ± 0.025 |

**Table S6** MD properties; averages and standard deviations over 3 MD replicas.

| Sequence | Bending persistence length (Å) | Torsional persistence length (Å) | Standard deviation of bending angle (degrees) |
|---|---|---|---|
| 1TA | 316 ± 54 | 617 ± 31 | 12.2 ± 1.2 |
| 4TA | 406 ± 72 | 670 ± 80 | 10.6 ± 1.6 |
| 2TA | 424 ± 55 | 633 ± 145 | 10.5 ± 1.2 |
| 4AT | 561 ± 72 | 742 ± 54 | 9.6 ± 0.9 |
| 1AT | 484 ± 87 | 722 ± 28 | 9.8 ± 1.2 |
| 3AT | 444 ± 18 | 654 ± 40 | 10.1 ± 0.7 |
| 4AA | 591 ± 71 | 712 ± 149 | 8.7 ± 0.1 |
| 4TT | 573 ± 47 | 698 ± 61 | 8.8 ± 0.2 |

**Figure S8** Step parameters for central $X_5U_6Y_7$ steps. Averages and standard deviations over all sequences and all simulations.

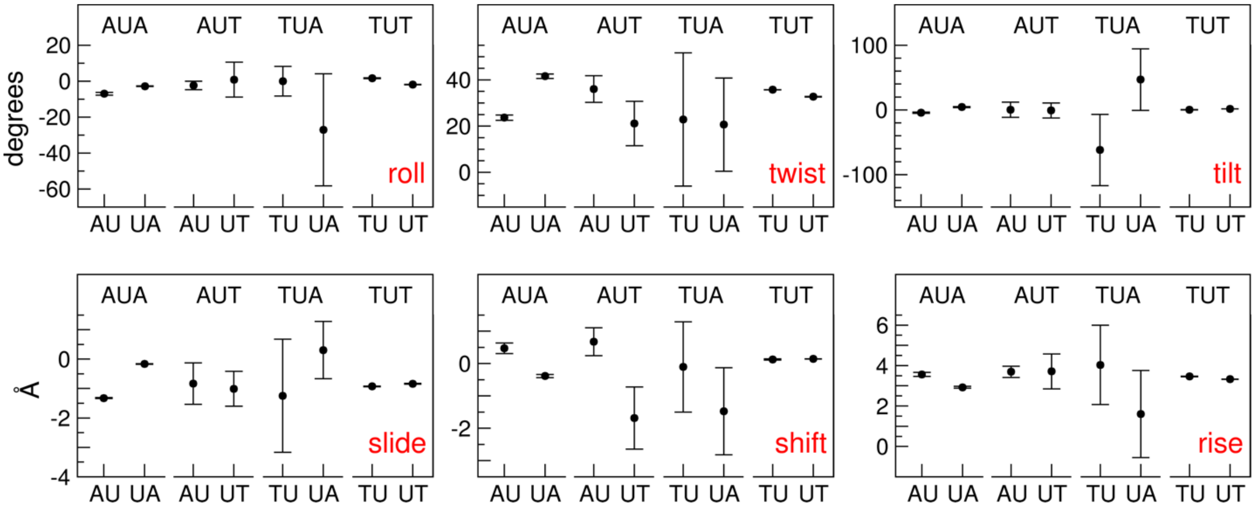

**Figure S9** Michaelis-Menten parameters ($K_m$ and $V_{max}$) from Table S2 plotted against $\alpha_0$ values from table S5. Results indicate that the values of $K$m, but not $k_{cat}$, are dictated by the mechanical properties of the substrate.

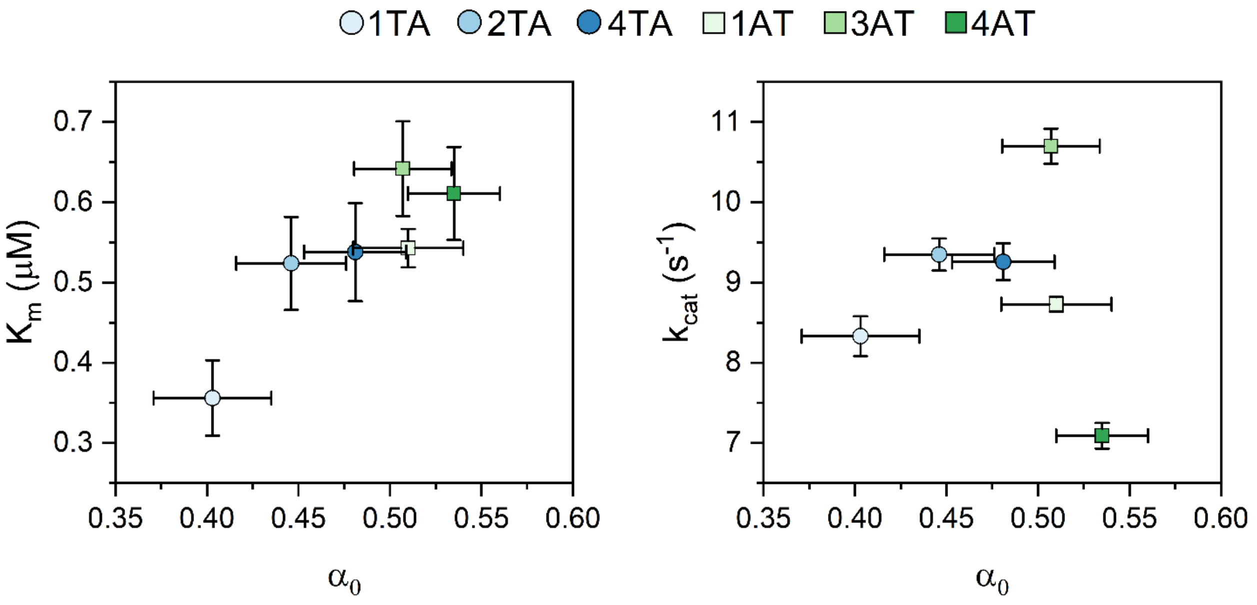


**Figure S10** Correlation of MD properties with $k_{ex}$. A) Bending persistence length (in Å), B) torsional persistence length (in Å), C) standard deviation of the bending angle (in degrees). Averages and standard deviations over 3 MD replicas; $k_{ex}$ in $s^{-1}$. Red lines show linear regression of all data with correlation coefficients of -0.889 (A), 0.739 (B), and 0.788 (C); blue lines linear regressions excluding the 4TA sequence with correlation coefficients of -0.504 (A), 0.054 (B), and 0.239 (C).

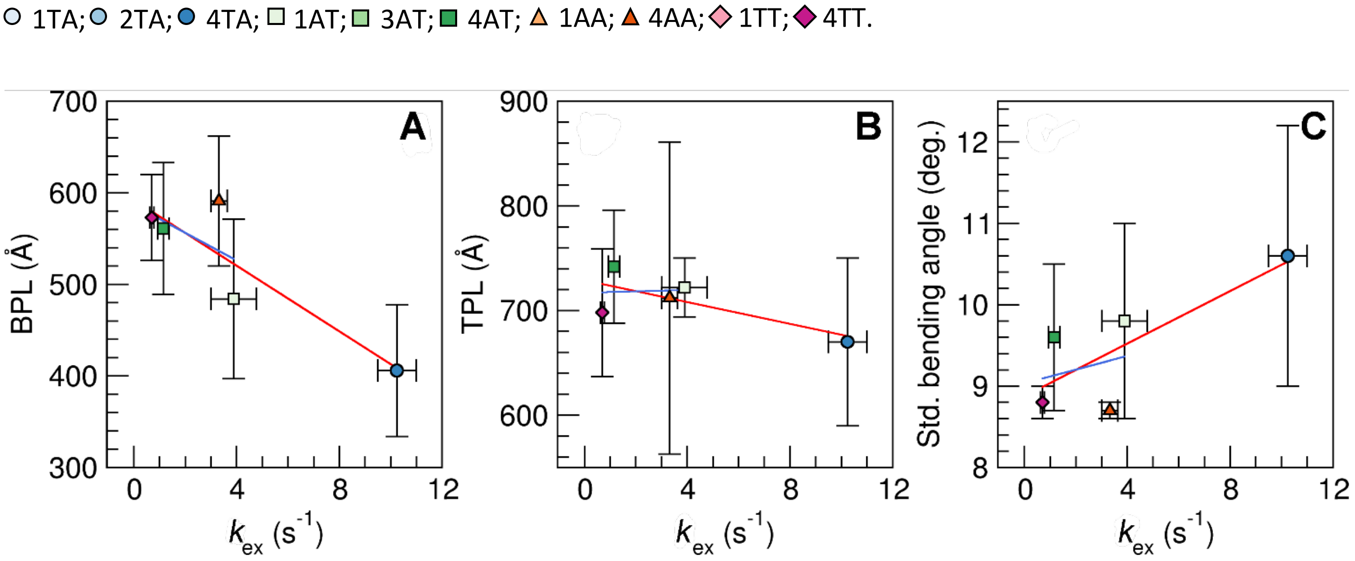
